# Frizzled-7 Identifies Platinum Tolerant Ovarian Cancer Cells Susceptible to Ferroptosis

**DOI:** 10.1101/2020.05.28.121590

**Authors:** Yinu Wang, Guangyuan Zhao, Salvatore Condello, Hao Huang, Horacio Cardenas, Edward Tanner, Jian-Jun Wei, Yanrong Ji, Junjie Li, Yuying Tan, Ramana Davuluri, Ji-Xin Cheng, Daniela Matei

## Abstract

Defining traits of platinum tolerant cancer cells could expose new treatment vulnerabilities. Here, new markers associated with platinum tolerant cells and tumors were identified by using *in vitro* and *in vivo* ovarian cancer (OC) models treated repetitively with carboplatin and validated in human specimens. Platinum-tolerant cells and tumors were found to be enriched in ALDH (+) cells, formed more spheroids, and expressed increased levels of stemness-related transcription factors compared to parental cells. Additionally, platinum-tolerant cells and tumors highly expressed the Wnt receptor, *Frizzled 7* (*FZD7*). FZD7 knock down improved sensitivity to platinum, decreased spheroid formation, and delayed tumor initiation. The molecular signature distinguishing FZD7(+) from FZD7(-) cells included *epithelial-to-mesenchymal (EMT), stemness*, and *oxidative phosphorylation* enriched gene sets. Overexpression of *FZD7* activated the oncogenic factor *Tp63*, driving upregulation of glutathione metabolism pathways, including glutathione peroxidase 4 (GPX4), which protects cells from chemotherapy-induced oxidative stress. FZD7(+) platinum-tolerant OC cells were more sensitive and underwent ferroptosis after treatment with GPX4 inhibitors. *FZD7, Tp63* and glutathione metabolism gene sets were strongly correlated in the OC Tumor Cancer Genome Atlas (TCGA) database and in human OC specimens residual after chemotherapy. These results support the existence of a platinum-tolerant cell population with partial stem cell features, characterized by FZD7 expression and dependent on FZD7-β-catenin-Tp63-GPX4 pathway for survival. The findings reveal a novel therapeutic vulnerability of platinum tolerant cancer cells and provide new insight into a potential “persister cancer cell” phenotype.

## Introduction

Ovarian cancer (OC) is the leading cause of death from female gynecological cancers. Although considered initially a highly chemo-responsive tumor, three quarters of patients with OC experience tumor relapse within two years of completing adjuvant platinum-taxane chemotherapy and recurrent, resistant OC is fatal (1). Understanding the underpinnings of chemo-resistance could lead to new therapies to eradicate such cells. “Persister” or drug-tolerant cells have been described as cells surviving cytotoxic drug exposure (2) and represent a reservoir for the outgrowth of drugresistant clones (3). Initially described as cells surviving exposure to targeted biological agents, the term is expanding to include chemotherapy-tolerant contexts. Recent studies in various cancers have reported the molecular signature on which “persister” cells depend, including upregulation of stemness factors, mesenchymal-like gene expression, enrichment in glutathione peroxidase 4 (GPX4) and of other genes related to lipid peroxidation, which antagonize ferroptosis allowing cells to survive after cytotoxic drug exposure (2, 4, 5). Small molecule inhibitors targeting GPX4 were shown to block lipid peroxidation and eliminate tyrosine kinase receptor inhibitor tolerant cells through ferroptosis (2). It has been suggested that “persister” cells share characteristics with cancer stem cells (CSCs), but also have unique traits, distinct from stemness. Specific markers to allow their identification and early targeting remain elusive. Here we sought out to characterize ovarian cancer cell phenotypes linked to platinum tolerance.

Our and other previous studies showed that while chemotherapy is effective at cytoreducing the mass of heterogenous cancer cells, residual tumors persist and are enriched in CSCs (6, 7). Ovarian CSCs share some of the normal stem cells’ characteristics, including the ability to self-renew, differentiate, express specific stem cell surface markers (6, 8, 9), and exhibit greatly enhanced tumor initiation capacity (TIC) (10). Importantly, ovarian CSCs possess a phenotype associated with drug resistance, including enhanced DNA repair, diminished apoptotic responses, increased efflux mechanisms and enhanced antioxidation defense (8, 11), which allow them to escape from chemotherapy. The boundaries between stemness and chemotherapy tolerant phenotypes remain blurry and while an overlap exists, it is assumed that distinct pathways drive the two entities.

As platinum tolerant cancer cells drive tumor relapse, decreasing survival rate of women with OC, we aimed to identify specific markers, by using *in vitro* and *in vivo* models of repeated exposure to the cytotoxic agent. We observed that platinum-tolerant cells and tumors contained an increased ALDH+ cell population, expressing stemness related transcription factors (TFs), and able to form more spheroids compared to chemotherapy naïve cells. We identified the *Frizzled 7* receptor *(FZD7)* as a novel cell surface marker significantly upregulated in the platinum tolerant cell population. *FZD7* knock down increased sensitivity to platinum, decreased spheroid formation *in vitro,* and delayed tumor initiation *in vivo*. FZD7(+) cells harbored a “persister cell”-like molecular signature, including down-regulated genes associated to *DNA damage response*, upregulated *epithelial to mesenchymal* (*EMT) and stemness* associated transcripts, and decreased expression of genes associated with *oxidative phosphorylation*. Expression of the antioxidant enzyme, GPX4 was increased in FZD7(+) platinum-tolerant cells, rendering these cells sensitive to treatment with GPX4 inhibitors. Mechanistically, *FZD7* expression caused activation of the transcriptional regulator, *Tp63*, which drove upregulation of glutathione metabolism genes, protecting cells from oxidative stress. In all, our results support the existence of a “persister” cell population, tolerant to platinum, sharing traits with CSCs, marked by upregulation of the receptor *FZD7* and harboring a dependency on *FZD7-b catenin-Tp63* mediated GPX4 expression and anti-oxidant activity.

## Materials and methods

### Human specimens

Deidentified high grade serous ovarian tumors (HGSOC) and associated malignant ascites were collected and processed fresh from consenting patients (Northwestern University IRB#: STU00202468). Tumor tissues were enzymatically disassociated into single cell suspensions and cultured as previously described (12). A tissue microarray (TMA) was built from deidentified HGSOC specimens (n=23) from patients who had undergone 3-6 cycles of platinum-taxane neoadjuvant chemotherapy (IRB approved CSR protocol #1247). Each specimen was entered in duplicate and fallopian tube epithelium (n=6) served as controls. Patients’ characteristics are in Table S3.

### Cell lines and culture conditions

SKOV3 and OVCAR3 cells were purchased from the American Type Culture Collection (ATCC). OVCAR5 cells were a generous gift from Dr. Marcus Peter, Northwestern University, COV362 cells were from Dr. Kenneth Nephew, Indiana University, immortalized human fallopian tube luminal epithelial cells (FT190) were from Dr. R. Drapkin of University of Pennsylvania (13). PE01 and PEO4 cells were from Sigma Aldrich. Cell culture conditions are in Supplemental Material (SM).

### Chemicals and reagents

RSL3 was purchased from Fisher Scientific (Cat# 611810). ML210 (Cat# SML0521), cisplatin (Cat# 1134357), and carboplatin (Cat# C2538) were from Sigma-Aldrich.

### *In vitro* development of platinum resistant cells

To generate platinum resistant OC cells *in vitro,* SKOV3, OVCAR5, COV362, and OVCAR3 cells were treated with 3 or 4 repeated or increasing doses of cisplatin or carboplatin for 24 hours. Surviving cells were allowed to recover for 3 to 4 weeks before receiving the next treatment. Changes in resistance to platinum were estimated by calculating half maximal inhibitory concentration (IC_50_) values as described below.

### In vivo experiments

Animal studies were conducted according to a protocol (# IS00003060) approved by the Institutional Animal Care and Use Committee of Northwestern University. To develop platinum resistant OC cells *in vivo,* female (6-8 weeks old) athymic nude mice *(Foxn1^nu^,* Envigo) were injected subcutaneously (s.c.) with 2 million SKOV3 or OVCAR3 cells, or intraperitoneally (i.p.) with 2 million OVCAR5 cells to induce tumors. Mice were treated i.p. with PBS (control) or 25 mg/kg carboplatin (n = 3-5), once-a-week for 3 weeks starting when xenografts were > 10 mm^3^ (SKOV3 and OVCAR3) or 2 weeks after i.p. inoculation (OVCAR5). Length (l), width (w) and height (h) of sc xenografts were measured weekly using digital calipers and tumor volume (v) was calculated as v = ½ × l × w × h. Tumors were collected 1 week after last treatment and were used for isolation of cancer cells, RNA and protein extraction, or fixed with 10% formalin neutralizing buffer (Formal-fix, Thermo Scientific, Ref# 9990244) for IHC. Two million OVCAR5 cells transduced with shRNAs targeting *FZD7* (OVCAR5_shFZD7) or scrambled shRNA control (OVCAR5_shctrl, n = 10 per group) were injected sc in 6-8 weeks female nude mice. Time to tumor initiation and tumor growth was monitored, as described above. Experiments using PDX tumors were performed in the Developmental Therapeutics Core (DTC) of the Lurie Cancer Center, as previously described (14) and following a similar protocol (see SM).

### Isolation of tumor cells

Tumors from patients or xenografts were minced and enzymatically dissociated in Dulbecco’s modified Eagle’s medium/F12 (Thermo Fisher Scientific, Ref# 11320) containing collagenase (300 IU/ml, Sigma-Aldrich, Cat#C7657) and hyaluronidase (300 IU/ml, Sigma-Aldrich, Cat# H3506) for 2-4 hours at 37°C. The tissue digest was passed several times through a 16-18G needle using Cell Stripper (Corning, Cat# 25-056-CI) to dissociate remaining cell aggregates. Red blood cell lysis used RBC lysis buffer (BioLegend, Cat#420301), followed by DNase (Qiagen, Cat# 79254) treatment and filtering through a 40μm cell strainer (Fisher Scientific, Cat#NC0147038) to yield single cells suspension.

### Aldefluor assay and flow cytometry

Aldehyde dehydrogenase (ALDH) activity was measured using an Aldefluor assay kit (Stemcell Technologies, Cat#01700, Cambridge, MA, USA) following the manufacturer’s instructions and as described previously (6)

### Cell survival assay

Cell survival was measured with a Cell Counting Kit 8 (CCK8, Dojindo Molecular Technologies, Cat# CK04, Rockville, MD, USA), following the manufacturer’s protocol. Absorbances (450 nm) were measured with a microplate reader (BioTek ELX800, BioTeK, Winooski, VT).

**Spheroid formation and clonogenic assays** were performed as described previously (see SM)(15, 16).

### Extreme limited dilution assay

A serial dilution of OVCAR5_shControl or OVCAR5_shFZD7 cells (5, 10, 50, 100, 500, 1000, and 5000 cells) were sorted by FACS directly into 96-well low-attached plates and cultured in MammoCult medium for 14 days as described above. Each dilution included 10 replicates. The total number of wells containing spheroids for each dilution were counted. The CSC frequency and statistical significance were determined using ELDA software at http://bioinf.wehi.edu.au/software/elda/(17).

### Half maximal inhibitory concentration (IC_50_)

The IC_50_ values for the various treatment compounds were determined by the CCK8 assay as described in SM. IC_50_ values were determined by logarithm-normalized sigmoidal dose curve fitting using Prism 6 software (GraphPad Software Inc., San Diego, CA).

### Lipid peroxidation assay

Intracellular lipid peroxidation was determined by a Lipid Peroxidation Assay (Sigma-Aldrich, Cat# MAK085) following the manufacturer’s protocol. Briefly, pellets of 1 or 3 million cells were homogenized on ice in 300 μl of the malondialdehyde (MDA) lysis buffer containing 3μl of butylated hydroxytoluene (BHT). After removal of insoluble materials, 600 μl of thiobarbituric acid (TBA) solution was added into each sample and incubated at 95°C for 60 minutes. Fluorescence was measure at 532/553 nm (excitation/emission) by using a microplate reader (SpectraMax GeminiXS, Molecular Devices).

### Cancer stem cell array

Human cancer stem cells RT^2^ Profiler PCR arrays were purchased from Qiagen (Cat# PAHS-176ZA-12) and used in combination with a 7900 HT real-time PCR system (Applied Biosystems). Data were normalized using the housekeeping genes included in the array, and relative mRNA amounts were calculated employing the 2^-ΔΔCt^ method. was Analyses were performed using a Microsoft Excel algorithm provided by the manufacturer and as described previously (15).

**Cell transfection, RNA extraction, quantitative RT-PCR analysis, western blotting and IHC** were performed according to previously published methods (15) described in SM.

### Oxygen consumption rate

Cells were seeded on 96-well plates at100000 cells/well and incubated overnight. Ten μL of extracellular O2 consumption reagent (Oxygen Consumption Rate Assay kit, Abcam Cat#197243) were added to each well, and fluorescence was measured with a plate reader (SpectraMax i3X, Molecular Devices, San Jose, CA, USA) at 3 min intervals for 180 min at excitation/emission = 380/650 nm. Alternatively, oxygen consumption was measured using a Seahorse assay. Briefly, OVCAR5 shctrl and shFZD7 cell lines were seeded in Seahorse 96-well microplate (Agilent, Cat#102416-100, Santa Clara, CA, USA) at a density of 10-80K per well.

After incubation overnight, oxygen consumption was measured and calculated by Seahorse XFe96 Analyzer (Agilent, Santa Clara, CA, USA).

### ODIPY staining for lipid peroxidation

Cells were treated as described in SM. After treatment, cells were stained with BODIPY 581/591 C11 (5 μM) for an hour at 37°C, washed with PBS, and fixed with 4% PFA on ice for 30 mins. The mean fluorescence intensity (minimum of 10,000 events per condition) was measured by FACS (LSR Fortessa, BD, Franklin lake, NJ). BODIPY emission was recorded on channels for FITC at 520nm and PE at 580nm. Data were displayed as histograms and mean fluorescence intensity of FITC was calculated.

### Intracellular reactive oxygen species (ROS)

Intracellular ROS levels were measured by monitoring the oxidation of cell permeable 2′,7′-dichlorofluorescein diacetate (DCFHDA, Sigma-Aldrich) to fluoros-pectrophotomete at excitation and emission wavelengths of 480 and 535 nm, respectively, measuring intracellular hydroxyl, peroxyl and other ROS activity, according to the manufacturer’s instructions (see SM).

### RNA sequencing (RNA-seq) and data analysis

The RNA-seq libraries (n=3 per experimental group) were prepared using the NEBNext Ultra II RNA library prep kit from Illumina (New England Biolabs Inc., Ipswich, MA, see SM). Differentially expressed genes were determined by exact test analysis followed by multiple hypothesis correction using false discovery rate (FDR) on the edgeR software (18). Genes with FDR ≤ 0.05 were considered differentially expressed. Normalized counts for all genes were ranked and subject to Gene Set Enrichment Analysis (19). Data are deposited in GEO (GSE148003; token: olwhsawwtbuhjuf).

### Analysis of data from The Cancer Genome Atlas (TCGA)

Processed Transcript-Per-Million (TPM) data were downloaded from TCGA-TARGET-GTEx Toil RNA-Seq Recompute Compendium (20) on UCSC Xena browser (21). 419 ovarian cancer samples from TCGA and 88 healthy ovarian control samples from GTEx were obtained. For isoform-level analysis, lowly expressed transcript isoforms were filtered out if they were not expressed in ≥ 90% of all samples. All genes and isoforms were annotated using Ensembl 82 (GRCh38.p3). **Correlation analysis between gene pairs:** Pearson correlation coefficient was calculated using log-transformed foldchange values of two genes of comparison. Outliers and influential points, with leverage exceeding twice of average of diagonal elements of hat matrix and Cook’s distance exceeding 10% confidence ellipsoid, were removed. The test statistic follows a t-distribution with n-2 degrees of freedom and the statistical significance of the correlation was determined by corresponding p-value. **Survival analysis:** Kaplan-Meier survival curves were plotted using R package ‘survival’ (R version 3.6.0). The high or low expression groups were defined based on statistically determined cutoff point that maximizes absolute value of the standardized two-sample linear rank statistic (22). The statistical significance of survival difference between groups with high/low level of expression was determined using log-rank test.

### Statistical analyses of experimental data

All data are presented as means values ± SD of triplicate measurements. Two-tailed Student’s *t*-test or ANOVA (one-way or two-way) were used to determine effects of treatments. P < 0.05 were considered significant. All analyses were performed using Prism 6.0 software (GraphPad Software).

## Results

### Stemness and ferroptosis signatures are enriched in platinum-tolerant cancer cells

Platinum resistant OC cells were generated through repeated *in vitro* exposure of OC cell lines (OVCAR3, OVCAR5, COV362 and SKOV3) to platinum at IC_50_ concentrations (Supplemental Figure S1A), while platinum resistant xenografts were obtained by treating tumor harboring mice with carboplatin for 4-6 weekly cycles (Figure S1B). Repeated platinum exposure of OC cells induced a stable phenotype, with a least 2-fold increase in platinum IC_50_ (Supplementary Table S1) compared to parental chemotherapy naïve cells. Platinum tolerant OC cells were enriched in ALDH (+) cells (Figure 1A and Figure S2A), formed increased numbers of spheroids (Figure 1B and Figure S2B), and contained cells expressing CSC-related TFs (*Oct4, Nanog* and *Sox 2*) (Figure 1C and Figs. S2C-D) compared to controls. RNA sequencing compared platinum-tolerant vs. parental naive cells, with transcriptomic signatures revealing enrichment in *stemness* associated gene sets (Figure 1D and Fig. S2E). Similar observations were made in *in vivo* models, including carboplatin treated OVCAR3 (Figs. 1E-F) and SKOV3 (Figure S2F) intraperitoneal xenografts. ALDH(+) cells were found to be enriched in the carboplatin treated xenografts (Figure 1E) compared to PBS-treated tumors and stemness associated genes (*ALDH1A1, Oct4, Nanog* and *Sox 2*; Figure 1F and Figure S2F) were upregulated in xenografts after carboplatin treatment compared to controls, as we noted previously (6, 23).

**Figure 1.**
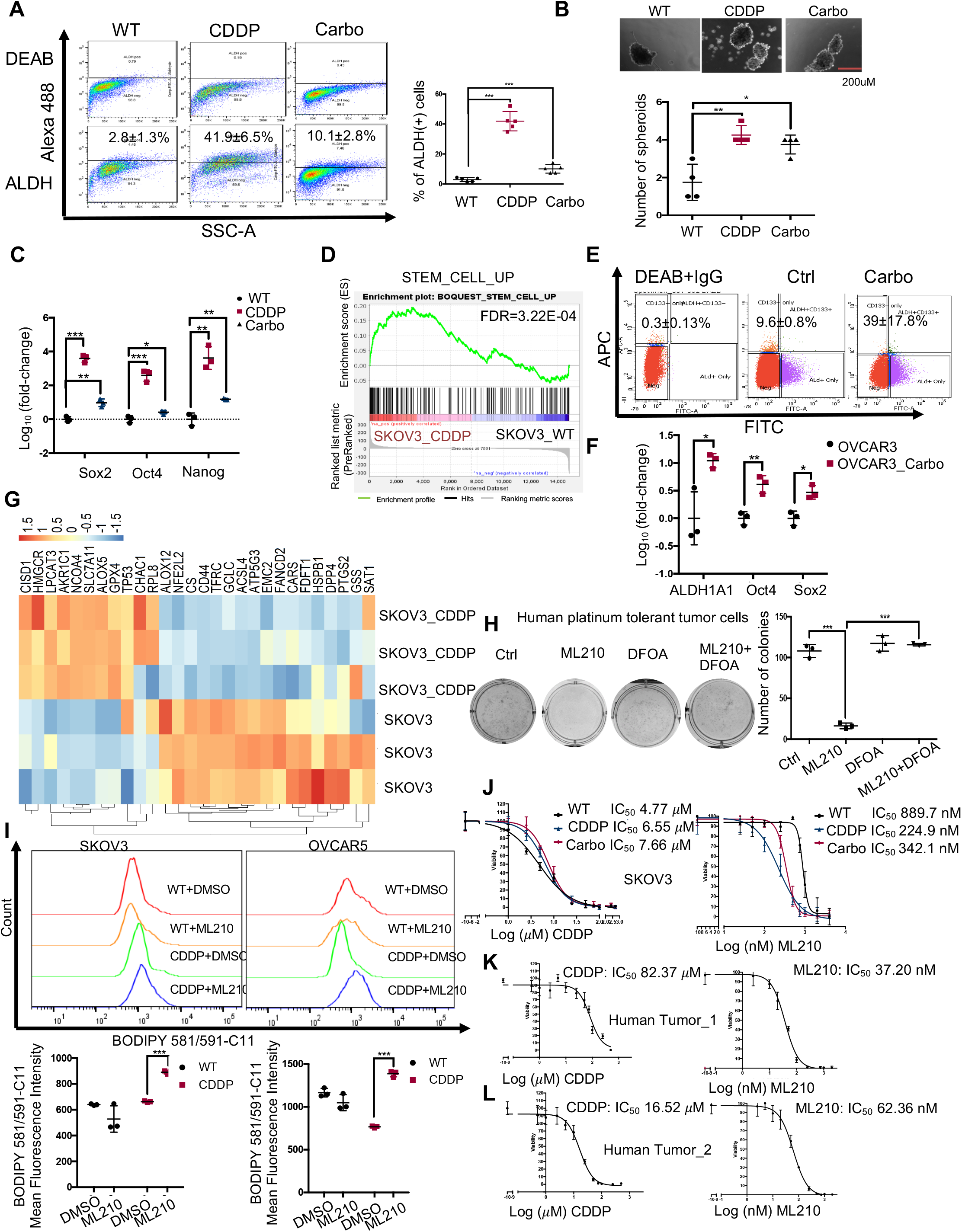
Stemness and ferroptosis signatures are enriched in platinum tolerant OC cells. **(A)** Representative FACS side scatter analysis of the ALDH(+) population (left), and percentage (mean ± SD, n=5) of ALDH(+) cells (right) in parental (WT), cisplatin tolerant (CDDP), and carboplatin tolerant (Carbo) SKOV3 cells. **(B)** Representative images (top), and numbers (mean ± SD, n=4) of spheroids (bottom) formed by 1000 parental (WT), cisplatin tolerant (CDDP), and carboplatin tolerant (Carbo) SKOV3 cells after 7 days of culture under non-attachment conditions. **(C)** mRNA levels (fold-change ± SD, n=3) of stemness-related transcription factors (*Sox2*, *Oct4* and *Nanog*) measured by real-time RTPCR in cisplatin (CCDP) and carboplatin tolerant (Carbo) SKOV3 cells compared with parental SKOV3 cells (WT). **(D)** GSEA shows upregulated stemness pathway (Stem Cell_UP) in SKOV3 CDDP-tolerant vs. parental cells (FDR=3.22E-04). **(E, F)** FACS side scatter analysis of percentage of ALDH(+) cells (E), and fold-change (mean ± SD, n=3) *ALDH1A1*, *Nanog*, and *Oct4* mRNA expression levels measured by real-time RT-PCR (F) in OC s.c. xenografts from mice injected with OVCAR3 cells and treated with PBS (Ctrl) or carboplatin (Carbo) to induce platinum resistance. **(G)** Hierarchical clustering heatmap for differentially expressed (FDR<0.05) ferroptosis-related genes in cisplatin tolerant (SKOV3_CDDP) vs. control SKOV3 cells. Gene expression was measured by RNA seq (n=3 replicates/group). **(H)** Representative pictures of a colony formation assay (left), and numbers (mean ± SD, n=3) of colonies (right) developed after 14 days from culturing 4000 cells isolated from platinum resistant primary HGSOC tumors treated with DMSO (Ctrl), the GPX4 inhibitor ML210 (500nM), the iron chelator DFOA (800nM), or ML210 plus DFOA combination for 24 hours. **(I)** Fluorescence histograms (top), and mean (± SD, n=3) fluorescence (bottom) of BODIPY 581/591-C11 staining show lipid peroxidation in SKOV3 and OVCAR5 parental (WT) cells and cisplatin tolerant (CDDP) cells treated with DMSO or ML210 (1 uM) for 20 hours. **(J)** Cell survival curves of SKOV3 wild type (WT), cisplatin tolerant (CDDP) and carboplatin tolerant (Carbo) cells in response to treatment with cisplatin (left) or GPX4 inhibitor ML210 (right). The cisplatin and ML210 IC_50_ values are shown for each cell subline. **(K-L)** Cell survival curves of cancer cells isolated from primary HGSOC tumors and treated with cisplatin (left), or GPX4 inhibitor ML210 (right). The IC_50_ for cisplatin and ML210 are shown. For all comparisons: *P<0.05, **P<0.01, ***P<0.001.

Given the possibility that CSCs would be more resistant to chemotherapy due to upregulated anti-redox mechanisms (11) and considering a recently proposed association between oxidative stress and ferroptosis, a new form of cell death triggered by oxidized lipids, we examined a gene set related to “ferroptosis” (24) in platinum tolerant compared to parental OC cells. Clear differences including upregulated genes involved in glutathione metabolism and anti-oxidant defense mechanisms were observed in platinum-tolerant OC cells vs. chemotherapy-naïve cells (SKOV3, Fig. 1G and OVCAR5, Fig. S2G). The anti-oxidant sellenoprotein gluthatione peroxidase 4 (*GPX4*), was among the upregulated genes in platinum tolerant cells. GPX4 inhibitors impede antioxidant defense mechanisms and promote death of cells dependent on this pathway (2). Platinum tolerant OC cells were more sensitive to the GPX4 inhibitor, ML210, compared to control cells (SKOV3, Fig. S3A; OVCAR5, Fig. S3B; COV362, Fig. S3C). ML210 induced inhibition of colony formation was inhibited by the iron chelator deferoxamine (DFOA), consistent with induction of a ferroptosis phenotype (Figs. S3A-C). Furthermore, primary human OC cells derived from malignant ascites from patients with recurrent platinum-resistant OC, were found to be dependent on GPX4, as ML210 potently reduced colony formation in these cells (Fig. 1H). Inhibition of colony formation induced by ML210 was also blocked by DFOA.

ML210 caused increased oxidized membrane lipids levels, as measured by flow cytometry using the C11-BODIPY dye in CDDP-tolerant compared to naïve cells, supporting increased susceptibility to ferroptosis of platinum tolerant OC cells (SKOV3 and OVCAR5, Fig. 1I; COV362, Fig S3D). Additionally, SKOV3 cisplatin and carboplatin-tolerant cells were more sensitive to ML210 compared to parental cells (IC_50_ of 224 nM and 342 nM vs. 889 nM, Figures 1I-J). Primary OC cells derived from malignant ascites associated with recurrent, platinum resistant OC, displayed resistance to platinum *in vitro* (Figures 1K-L, left panels; IC_50_ of 82μM and 16.52μM), and responded to low doses of the GPX4 inhibitors, ML210 (Figures 1K-L, right panels; IC_50_ of 37.20 nM and 62.36 nM) and RSL-3 (Figure S3E-F, IC_50_ of 14.20 nM and 17.72 nM). These results derived from multiple *in vitro* and *in vivo* OC models, including primary human cancer cells, support the existence of a “persister cell” phenotype induced by prior exposure to and development of tolerance to platinum, sharing partial stemness characteristics, and highly susceptible to ferroptosis.

### Frizzled 7 (FZD7) is upregulated in platinum tolerant OC cells and tumors

To identify potential markers linked to this “persister” phenotype, an RT-PCR based platform representing 90 cancer stemness associated genes was used. A number of known CSC markers were found to be upregulated in the platinum-tolerant cells (*CD44, PROM1, SOX2*), along with membrane transporters known to be associated with platinum-resistance (*ABCG2, ABCB5*), and regulators of EMT (*TGFBR1, SNAI1, BMP7, TWIST 1* and *2, SNAI 1* and *2,* see Table S2). Among transcripts representing membrane proteins, which could potentially be used as novel markers, *FZD7*, a transmembrane receptor involved in canonical Wnt/β-catenin/TCF and non-canonical Wnt/planar cell polarity (PCP) signaling (25, 26), was one of the top highly expressed transcripts (> 8-fold) in platinum tolerant compared to control cells (Supplementary Table S2 and Figure S4A). Increased *FZD7* expression levels were confirmed in platinum tolerant OC cell models (SKOV3, OVCAR5, and OVCAR3) generated as described, compared to control cells at *mRNA* (Figure 2A) and protein level (Figure 2B), but also in platinum resistant PEO4 cells compared to sensitive PEO1 OC cells, an isogenic cell line pair, derived from the same patient at different times during the disease course (PEO1 during a platinum sensitive recurrence and PEO4 during a platinum-resistant recurrence (27) (Figs. 2C-D). Likewise, *FZD7 mRNA* expression levels were upregulated in platinum treated SKOV3 (Fig. 2E) and OVCAR3 (Fig. 2F)-derived xenografts compared to vehicle-treated tumors.

**Figure 2.**
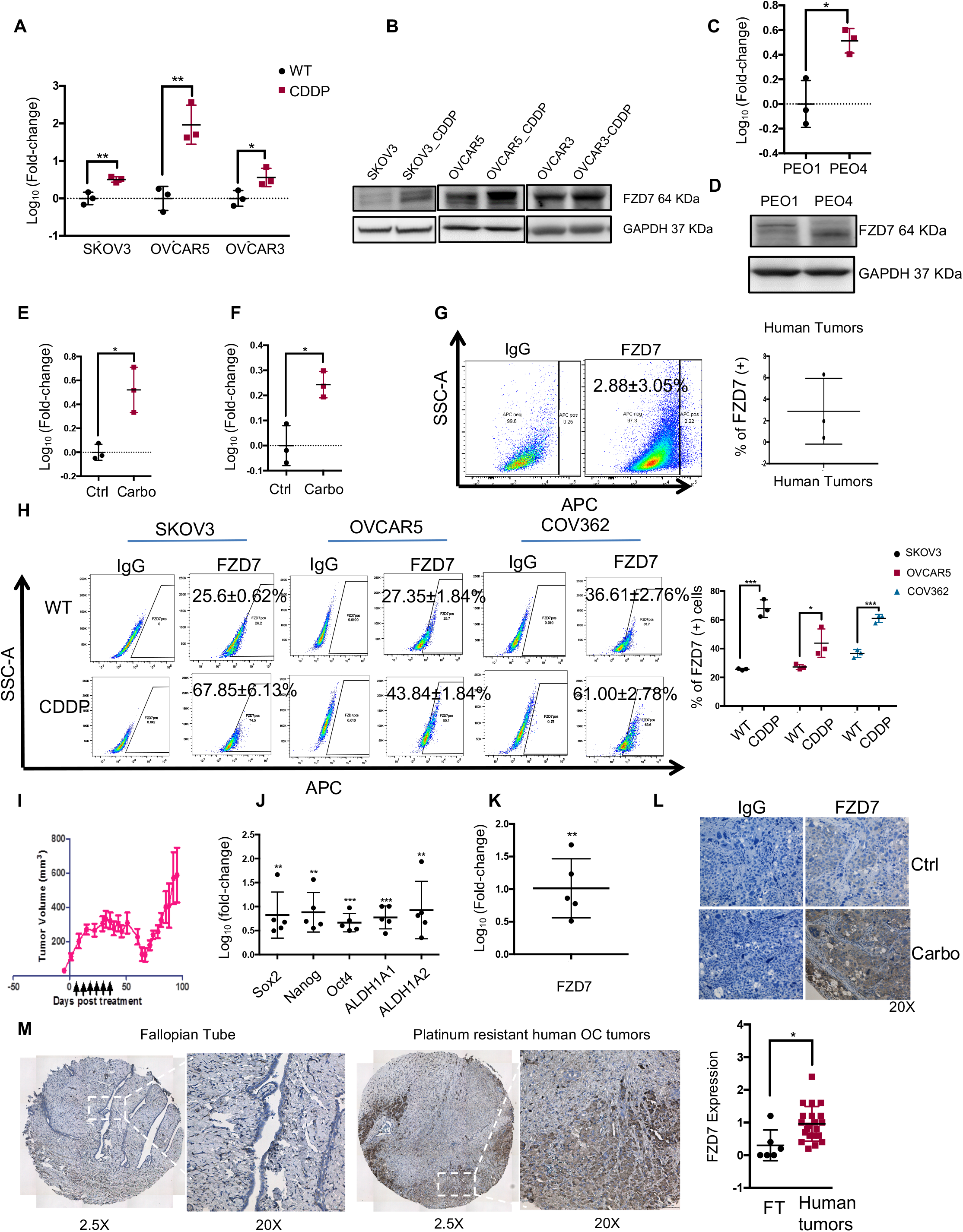
Frizzled 7 (FZD7) is upregulated in platinum resistant OCs. **(A)** Fold-change (mean ± SD, n=3) of *FZD7 mRNA* expression levels measured by real-time RT-PCR in cisplatin tolerant (CDDP) SKOV3, OVCAR5 and OVCAR3 OC cells compared with parental cell lines. **(B)** Western blotting shows FZD7 protein levels in SKOV3, OVCAR5 and OVCAR3 OC cells and their cisplatin tolerant (-CDDP) variants (n = 2). **(C, D)** *FZD7 mRNA* expression levels (mean foldchange ± SD, n = 3) measured by real-time RT-PCR (C), and FZD7 protein levels measured by western blotting (n = 2) (D) in platinum resistant PEO4 compared with platinum sensitive PEO1 OC cells. **(E)** Real-time RT-PCR measures fold-change (mean ± SD) of *FZD7 mRNA* levels in carboplatin-treated (Carbo) and control (Ctrl) SKOV3 (E) and OVCAR3 (F) subcutaneous xenograft tumors (n = 3 animals per group). **(G)** Representative FACS side scatter analysis (left) and average (±SD, n = 3) of FDZ7(+) cells dissociated from HGSOC tumors. **(H)** FACS side scatter analysis of FDZ7(+) cells (left), and percentage (mean ± SD, n=3) of FZD7(+) cells (right) in SKOV3, OVCAR5 and COV362 parental (WT) cells and their cisplatin tolerant (CDDP) sublines. **(I)** Development (mean volume ± SD, n = 5) of subcutaneous HGSOC PDX tumors in NSG mice receiving carboplatin treatment (15mg/kg, weekly) to induce platinum tolerance. Arrows indicate the days of carboplatin treatment. Tumors were collected after residual tumors redeveloped. **(J)** *Sox2, Nanog, Oct4, ALDH1A1* and *ALDH1A2* mRNA expression levels (mean fold-change ± SD, n = 5) measured by real-time RT-PCR in carboplatin tolerant PDX xenografts compared to control PDX tumors generated as described in I. **(K,L)** Real-time RT-PCR measurements (mean fold-change ± SD, n=5) of *FZD7* mRNA levels (K), and representative images of FZD7 IHC staining (L) in carboplatin tolerant PDX tumors (Carbo) compared with control PDX tumors (Ctrl) generated as described in I. Magnification 20X. **(M)** Representative pictures of FZD7 IHC staining (left and middle) and FZD7 H-scores (mean ± SD) (right) in sections of normal fallopian tube (n=6) and platinum resistant HGSOC tumors (n=21) included in a tissue microarray. Pictures were taken at the indicated magnifications. For all indicated comparisons: *P< 0.05, **P<0.01, and ***P<0.001.

Flow cytometry was used to determine whether a FZD7 high (FZD7+) cell population is detectable in OC models. FZD7(+) cells were detected among cells dissociated from primary human ovarian tumors, previously untreated with chemotherapy, and represented ~ 3% of all cells (Figure 2G). In OC cell lines, FZD7(+) cells were identified as a distinct sub-population representing ~25-35% of cells (SKOV3, OVCAR5 and COV362; Figure 2H). Additionally, the FZD7(+) cell population was detectable and was enriched in CDDP-tolerant compared to parental cells (SKOV3, OVCAR5, COV362; Figure 2H), suggesting that this cell membrane receptor may be a marker for “persister” cells, pre-existing in the un-selected heterogeneous cell populations prior to exposure to platinum, and enriched after exposure to the cytotoxic drug.

Further, we used patient derived xenografts (PDX) generated from newly diagnosed high grade serous ovarian tumors implanted intra-bursally in the ovary of NSG mice (14). The original PDX tumors were subsequently passaged into second generation mice and treated weekly with carboplatin. After an initial response to platinum, recurrent tumors emerged (Figure 2I). Consistent with observations in OC cell lines and xenograft models, ALDH (+) cells were enriched in platinum tolerant PDX compared with PBS-treated tumors (Fig. S4B) and CSC-related transcription factors (*Sox 2, Nanog, Oct4),* as well as *ALDH1A1* and *ALDH1A2* were upregulated in platinum-tolerant PDX vs. control (Figure 2J). *FZD7* expression levels were also upregulated at *mRNA* (Figure 2K) and protein level (Figure 2L), as measured by immunohistochemistry (IHC) in platinum tolerant PDX vs. control, further supporting the concept that expression of this receptor is associated with a platinum-tolerant state.

Next, IHC assessed FZD7 expression levels in human HGSOC specimens collected after 3-6 cycles of neo-adjuvant chemotherapy, thus containing cells surviving after standard platinum-taxane chemotherapy, which are presumably chemotherapy-tolerant. Patients’ characteristics for these tumors are included in Supplementary Table S3. Increased FZD7 staining intensity (measured as H-score) was observed in cancer cells residual after chemotherapy in these specimens (n = 23), when compared to fallopian tube epithelium (control n = 6, Figure 2M, *p* = 0.02).

### Functional role of FZD7 in OC cells

To characterize the phenotype of FZD7(+) cells, fluorescence-activated cell sorting (FACS) separated cells expressing high endogenous levels of the receptor FZD7(+) vs. FZD7(-) cells from SKOV3, OVCAR5, COV362, and human OC tumors. Differences in *mRNA* expression levels between FZD7(+) and FZD7(-) cell are shown in Fig. 3A. FZD7(+) OC cells were less sensitive to cisplatin (CDDP) (Figures 3B-D; *p* < 0.05), supporting that the receptor marks a platinum-tolerant population. Additionally, FZD7(+) cells formed spheroids more efficiently compared to FZD7(-) cells (Figure 3E-F, *p* < 0.01), and expressed higher levels of stemness associated transcription factors, including *Nanog* and *Sox2* (SKOV3, Fig. 3G, OVCAR5, Fig. 3H; COV362, Fig. 3I; *p* < 0.05), supporting that they share stemness-related features.

**Figure 3.**
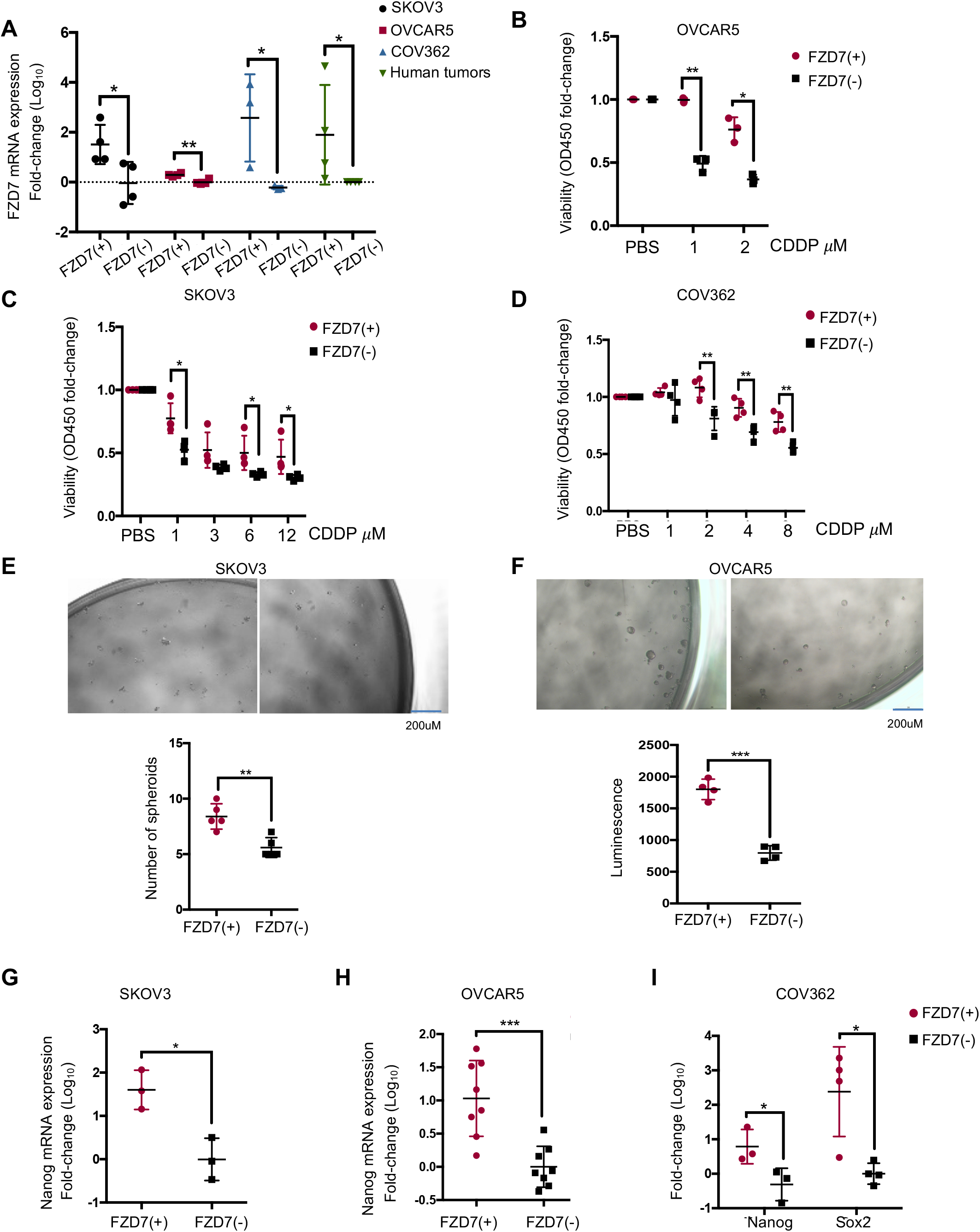
Functional roles of FZD7 in OC cells and molecular signature of FZD7(+) cancer cells. **(A)** Real-time RT-PCR measured *FZD7 mRNA* levels (mean fold-change ± SD, n=3-4) in FZD7 (+) and FZD7 (-) cells selected by FACS from SKOV3, OVCAR5, and COV362 cell lines and HGSOC tumors. **(B-D)** Cell viability (mean fold-change ± SD, n=4) of FZD7(+) and FZD7(-) cells selected by FACS from OVCAR5 (n=3) (B), SKOV3 (n=3) (C) or COV362 (n=4) (D) cells, plated, treated with the indicated doses of cisplatin (CDDP) for 24 hours, and cultured for additional three days. Cell viability was measured with the CCK8 assay. **(E, F)** Representative pictures and numbers (mean ± SD, n = 4-5) of spheroids formed after 7 days of culture by FZD7(+) and FZD7(-) cells sorted by FACS from SKOV3 (E) and OVCAR5 (F) OC cells. Spheroids were counted visually or their numbers (cells growing as spheres) were estimated by using a CellTiter-Glo 3D cell viability assay. **(G-I)** Real-time RT-PCR measures mRNA levels (fold-change ± SD) of the stem cell marker *Nanog* in FZD7(+) compared with FZD7(-) cells sorted by FACS from SKOV3 (J) (n=3) or OVCAR5 (I) (n=8) cells. **(L)** Real-time RT-PCR measured *mRNA* levels (fold-change ± SD, n=3-4) of the stem cell markers *Nanog* and *Sox2* in FZD7(+) compared with FZD7(-) cells sorted by FACS from COV362 OC cells. For all indicated comparisons: *P<0.05, **P<0.01, ***P<0.001.

To further examine *FZD7* functions, the receptor was knocked-down (KD) by stable transduction of shRNA or was transiently overexpressed in several OC cell lines. Decreased *FZD7 mRNA* expression was confirmed by Q-RT-PCR in SKOV3 and OVCAR5 cells transfected with two shRNA sequences targeting the receptor (Figs. 4A-B). *FZD7* knock down decreased spheroid formation (Figs 4A-B) and expression of stemness associated TFs (*Sox2* and *Nanog*; Figure 4C and Fig. S5A). *In vitro* serial limited dilution assay showed that receptor knock down in OVCAR5 cells caused a decreased stem cells frequency, as calculated by the ELDA software (28) in sh-control transfected cells vs. sh-FZD7 transduced cells, (p = 0.034, Fig. S5B). Likewise, *FZD7* KD in platinum-tolerant SKOV3_CDDP cells (Fig. S5C) decreased sphere formation (Fig. S5D). Stable *FZD7* KD in primary platinum resistant OC tumor cells caused decreased expression levels of the stemness-associated gene *ALDH1A1* (Figures 4D-E). Reversely, transient overexpression of *FZD7* in OC cells (Figs. 4E-F) promoted proliferation of SKOV3 and OVCAR5 cells as spheres (Fig. 4G) and increased expression of stemness associated TFs (*Sox 2* and *Nanog*, Fig. S5E-F). *FZD7* KD also decreased IC_50_ to cisplatin *in vitro* by ~ 2-fold (SKOV3, Fig. 4H; SKOV3_CDDP, Fig. 4I); while the receptor’s overexpression increased resistance to cisplatin (Fig. 4J).

**Figure 4.**
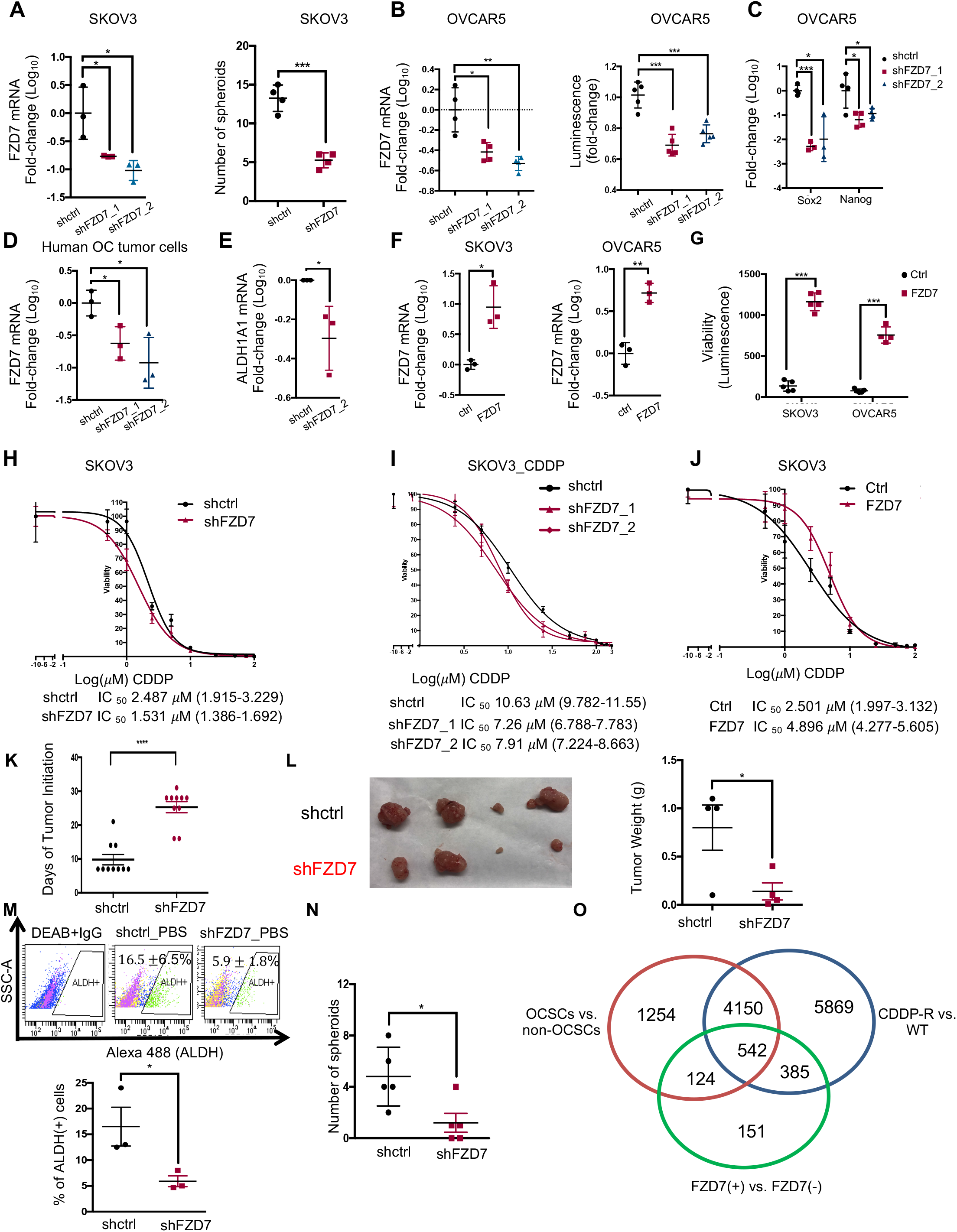
FZD7 regulates stemness characteristics of OC cells. **(A)** (Left) *FZD7 mRNA* levels (mean fold-change ± SD, n=3) measured by real-time RT-PCR in SKOV3 cells transduced with shRNAs targeting *FZD7* (shFZD7) cells vs. cells transfected with control shRNAs (shctrl). (Right) Numbers of spheroids (mean ± SD, n=4) formed by 2,000 shFZD7 or shctrl SKOV3 cells cultured for 14 days. Spheroids were counted under a microscope. **(B)** *FZD7 mRNA* expression levels (mean fold-change ± SD, n=4) determined by real-time RT-PCR (left), and numbers of spheroids assessed by measuring cell viability (mean fold change of luminescence ± SD, n=5) (right) in OVCAR5 cells transduced with shRNAs directed at *FZD7* (shFZD7) compared with cells transduced with control shRNAs (shctrl). Spheroids formation assay used 2,000 cells that were cultured for 7 days. Cell viability was determined with a CellTiter-Glo 3D assay. **(C)** *Sox2* and *Nanog* mRNA levels (mean fold-change ± SD, n=3-4) measured by RT-PCR in OVCAR5 cells transduced with shFZD7 vs. shctrl. Cells were generated as described in B. **(D)** *FZD7 mRNA* levels (mean fold-change ± SD, n=3) measured by real-time RT-PCR in platinum resistant primary HGSOC tumors cells transduced with shRNAs targeting *FZD7* (shFZD7) vs. cells transfected with control shRNAs (shctrl). **(E)** *ALDH1A1* mRNA levels (mean fold-change ± SD, n = 3) measured by RT-PCR in primary tumor cells transduced with shFZD7 cells vs. shctrl. **(F)** Average foldchange (± SD, n = 3) of *FZD7 mRNA* measured by real-time RT-PCR in SKOV3 (left) and OVCAR5 (right) cells transfected with FZD7-pcDNA3.1 vs. cells transfected with empty vector (ctrl). **(G)** Spheroid formation estimated with a CellTiter-Glo viability kit (bottom) from 1,000 ctrl and FZD7 expressing SKOV3 or OVCAR5 cells (described in F) and cultured for 7 days (n =4-5 per group). **(H-J)** Effects of cisplatin (CDDP) on cell survival measured by CCK8 assays in SKOV3_shctrl and SKOV3_shFZD7 (H), SKOV3 cisplatin tolerant cells (SKOV3_CDDP) transduced with shRNAs targeting FZD7 (shFZD7_1, _2) or control shRNA (I), and SKOV3 cells transfected with FZD7-pcDNA3.1 (FZD7) or empty vector (Ctrl) (J). Cells were treated with cisplatin for 24 hours and cultured for additional 3 days (n=3-4). The cisplatin IC_50_ value for each cell line variant is shown below the corresponding figure. **(K)** Days to initiation (mean ± SD, n = 10) of subcutaneous xenograft tumors induced by 2×10^6^ shctrl and shFZD7 OVCAR5 OC cells (generated as described in A) in nude mice. Time to initiation was the time from cancer cell inoculation to time when tumors were detectable (> 2mm in largest dimension). **(L)** Pictures and weights (mean ± SD, n=4) of tumor xenografts induced by OVCAR5_shctrl and OVCAR5_FZD7 cells. **(M)** FACS side scatter analysis of ALDH(+) cells (top), and percentage (mean ± SD, n=3) of ALDH(+) cells (bottom) in cell suspensions generated from OVCAR5_shctrl and OVCAR5_shFZD7 xenografts. **(N)** Numbers (mean ± SD, n=5) of spheroids (bottom) formed during 14 days by 1000 cells derived from cell suspensions generated from OVCAR5_shctrl or OVCAR5_FZD7 xenografts. **(O)** A Venn diagram shows the number overlapping and unique genes that differ in expression between OVCAR5-derived OCSCs (ALDH+CD133+) versus non-OCSCs (ALDH-CD133-), OVCAR5 cisplatin tolerant (CDDP-R) vs. parental (WT), and FZD7(+) versus FZD7(-) OVCAR5 OC cells. ALDH+CD133/ALDH-CD133- and FZD7+/FZD7-cells were sorted by FACS (see SM). *mRNA* levels were measured by RNAseq and statistical significance was set at FDR < 0.05. For all indicated comparisons: *P<0.05, **P<0.01, ***P<0.001.

To test the effects of FZD7 on tumor formation, a subcutaneous (s.c.) xenograft model was used. *FZD7* knock down in OVCAR5 cells delayed tumor initiation *in vivo* (sh-control 9.8±4.6 *days* vs. sh-FZD7 25.3±4.9 *days,* Figure 4K, *p* < 0.0001) and decreased tumor weight (0.80±0.41 *g* vs. 0.14±0.15 *g,p* = 0.04, Fig 4L, n = 4/group) and tumor size (Fig. S5G) *in vivo. FZD7* knock-down caused decreased ALDH(+) fraction in cell populations dissociated from xenografts (16.5±6.5% vs. 5.9±1.8%, *p* = 0.05; Fig. 4M, n = 3) and their spheroid forming ability (Fig. 4N, *p* = 0.02). Combined, these results support that FZD7 is linked to stemness and chemo-resistance in OC models.

### Molecular signatures of FZD7(+) cancer cells

To gain further insight into the functional role of FZD7 relative to stemness and chemo-responsiveness, gene signatures distinguishing FZD7(+) vs. FZD7(-) cells; ovarian CSCs (ALDH^+^CD133^+^) vs. non-CSCs (ALDH^-^CD133^-^), and CDPP-tolerant vs. platinum-naïve OVCAR5 cells were examined and integrated (Figure 4O). FZD7(+) and (-) were sorted by FACS (Fig. S5G). Ovarian CSCs and non-CSCs were FACS-sorted by using dual stem cell markers, CD133 and Aldefluor activity (Fig. S5I). There were 666 differentially expressed genes (DEG) shared between FZD7(+)/FZD7(-) and CSCs/non-CSCs datasets, 5404 DEGs being unique to CSCs and 536 DEGs unique to FZD7(+) cells (Figure 4O). Additionally, there were 927 DEGs overlapping between FZD7+/FZD7- and resistant/parental cells, and 10019 DEGs unique to the platinum-tolerant cells and 275 genes uniquely associated with FZD7(+) cells (Figure 4O). Overlapping DEGs between FZD7+/FZD7- and resistant/parental cells were enriched in *Cancer Stem Cell* and *DNA Repair* Signatures (Fig. S6A-B). Additionally, FDZ7(+) vs. FZD7(-) cells displayed signatures enriched in *stemness* (Fig. S6C), *EMT* (Fig. S6D) and *downregulated DNA damage response* genes (Fig. S6E). Together, these data suggest that FZD7(+) cells possess both shared, but also distinct features, relative to stemness and chemo-resistance, consistent with the phenotypes described above. Importantly, *mitochondrial and oxidative phosphorylation* gene sets were enriched among DEGs distinguishing FZD7(+) vs. FZD7(-) cells (Fig. S6F-H) and Ingenuity Pathway Analysis (IPA) identified *Oxidative Phosphorylation* and *Mitochondria Dysfunction* as the top enriched pathways in FZD7(+) cells, suggesting that the receptor marks a cell population harboring altered oxidative stress responses.

### FZD7 marks a cell population enriched in GPX4

Given that GPX4, an antioxidant enzyme which reduces reactive oxygen species (ROS), preventing formation of toxic lipid peroxides (29, 30), has been implicated in maintenance of normal mitochondrial function and oxidative phosphorylation (2, 29), we next examined whether *FZD7* expression impacted *GPX4* expression and function in platinum-tolerant OC cells. GPX4 expression levels were measured in FZD7(+) and FZD7(-) cancer cells and in platinum tolerant OC cells and tumors. *GPX4* levels were significantly increased in FACS sorted FZD7(+) vs. FZD7(-) cells derived from SKOV3, OVCAR5, COV362 cells or cancer cells dissociated from human tumors (Fig. 5A, *p* < 0.05). Furthermore, *FZD7* KD by shRNA in OVCAR5 and SKOV3 cells resulted in repressed *GPX4 mRNA* (Fig. 5B, *p* < 0.001, and Fig S7A, *p* < 0.05) and protein expression levels (Fig. 5C). GPX4 expression levels were also decreased in resistant SKOV3-CDDP cells (Fig. 5D) and in primary platinum resistant OC cells (Fig. 5E) transduced with shRNA targeting *FZD7.* Reversely, *FZD7* overexpression in SKOV3 and OVACR5 cells induced increased GPX4 *mRNA* and protein expression levels (Figs. 5F-H).

**Figure 5.**
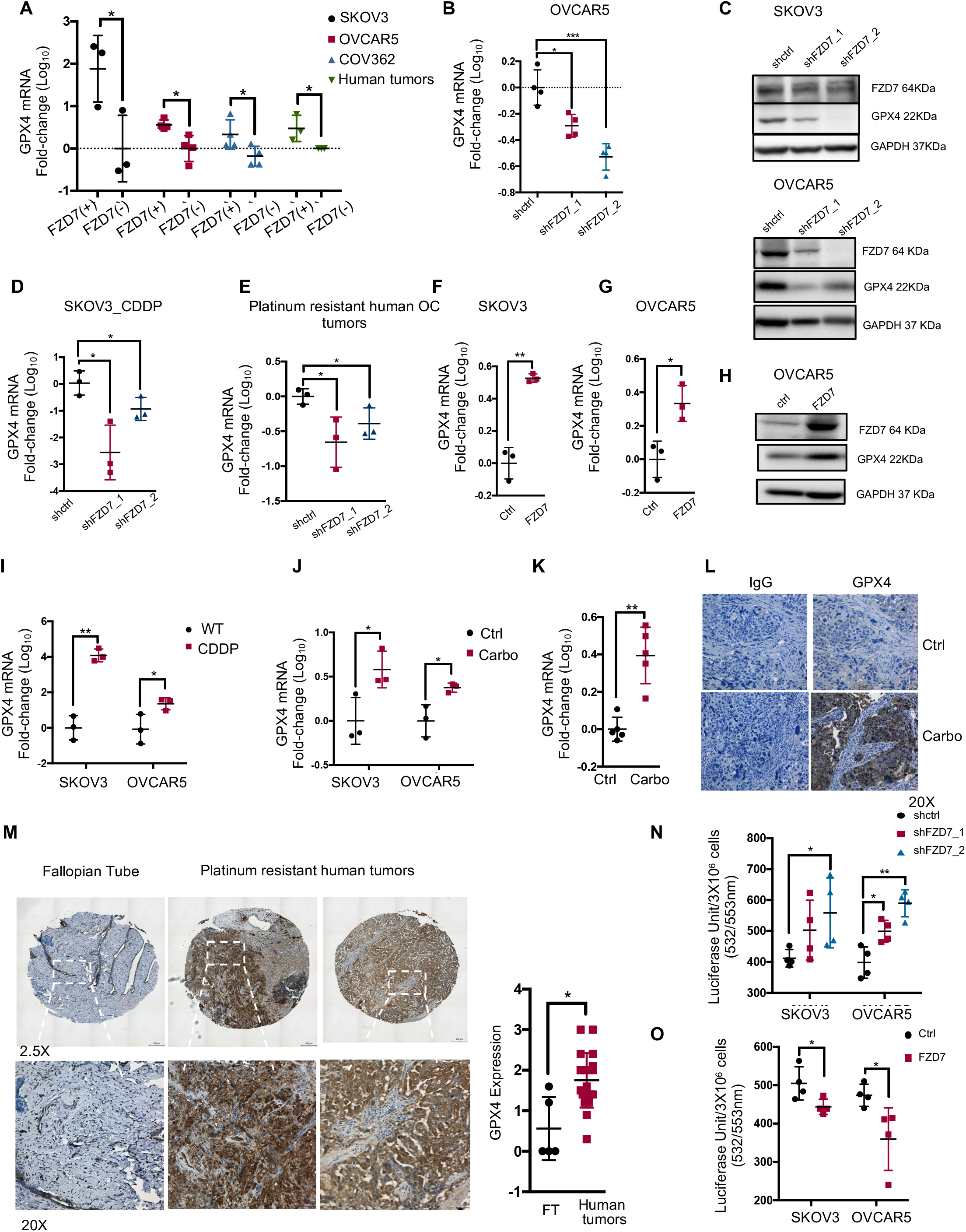
FZD7 regulates GPX4 and intracellular redox states. **(A)** Fold-change (mean ± SD, n=3-4) of *GPX4 mRNA* levels measured by real-time RT-PCR in FZD7(+) vs. FZD7(-) cells sorted by FACS from SKOV3, OVCAR5, COV362, OC cell lines and cell suspensions derived from HGSOC tumors. **(B)** Real-time RT-PCR measured fold-change (mean ± SD, n=4) of *GPX4 mRNA* expression levels in OVCAR5 cells transduced with shRNAs targeting *FZD7* (shFZD7) compared with cells transduced with control shRNAs (shctrl). **(C)** Western blotting for FZD7, GPX4 and GAPDH (loading control) in SKOV3 and OVCAR5 cells stably transduced with shctrl and shFZD7, as described in (B) (n = 2). **(D-E)** *GPX4 mRNA* expression levels (fold-change ± SD, n = 3) in SKOV3 cisplatin tolerant cells (CDDP) (D), and platinum resistant primary human HGSOC cells (E) transduced with shRNAs directed at *FZD7* (shFZD7) vs. cells transduced with control shRNAs (shctrl). **(F, G)** Average fold-change (± SD, n=3) of *GPX4 mRNA* levels in SKOV3 (F) and OVCAR5 (G) cells transfected with FZD7-pcDNA3.1 vs. cells transfected with empty vector (ctrl). **(H)** Western blotting of FZD7, GPX4 and GAPDH (loading control) in OVCAR5 cells transfected with FZD7-pcDNA3.1 compared with cells transfected with empty vector (ctrl) (n = 2). **(I-K)** Real-time RT-PCR measurements of *GPX4 mRNA* expression (mean fold-change ± SD) in SKOV3 and OVCAR5 cisplatin tolerant OC cells (CDDP) compared with parental lines (WT) (n = 3 per group) (I), SKOV3 and OVCAR5 xenografts from mice (n = 3 per group) treated with PBS (Ctrl) or carboplatin (25 mg/kg, once-a-week for 3 weeks) (J), and PDX tumors from mice receiving carboplatin (n = 5 per group) (K). **(L)** Representative images of GPX4 IHC staining in sections of control (Ctrl) and carboplatin tolerant PDX tumors. Magnification 20X. (**M)** Representative pictures of GPX4 IHC (left) and GPX4 expression assessed by H-scores (mean ± SD) (right) in sections of normal fallopian tube (n = 6) and chemoresistant ovarian carcinoma (n = 20). Pictures were taken at indicated magnification. Bottom pictures show inset areas at higher magnification (20X). **(N)** Intracellular lipid peroxidation measured with an MDA assay in SKOV3 and OVCAR5 cells transduced with scrambled shRNA (shctrl) or shRNAs targeting *FZD7* (shFZD7) and expressed as average luciferase units/3X10^6^ cells (± SD, n=4) **(O)** Intracellular lipid peroxidation measured with a MDA assay in SKOV3 and OVCAR5 cells transfected with FZD7 expression vector (FZD7) or control vector (Ctrl) and expressed as average luciferase units/3X10^6^ cells ± SD (n=4). For all indicated comparisons: *P<0.5, **P<0.01, ***P<0.001.

Furthermore, *GPX4* and *FZD7* mRNA expression levels were coordinately upregulated in platinum-tolerant OC models, including platinum resistant versus parental cells (SKOV3, OVCAR5, Fig. 5I, *p* < *0.05*), carboplatin-tolerant xenografts (Fig. 5J, *p* < *0.05*), and carboplatin-tolerant PDX tumors (Fig. 5K, *p* < 0.01). Increased *GPX4* expression in platinum tolerant ovarian PDX vs. control tumors was confirmed by IHC (Fig. 5L). To demonstrate the clinical significance of the findings, we used IHC for GPX4 in HGSOC specimens collected after neo-adjuvant chemotherapy. These specimens contain cancer cells residual after 3-6 cycles of treatment with platinum and taxanes. Significantly increased GPX4 staining was observed in these residual tumors (n = 23) when compared to fallopian tube epithelium, Fig. 5M, p = 0.02). Together, the results confirm a positive correlation between FZD7 and GPX4 expression in OC cells and in platinum-tolerant models, supporting that FZD7(+) cells have an increased anti-oxidant capacity.

To confirm that these correlations are functionally relevant, the enzymatic activity of GPX4 was measured by using the malondialdehyde (MDA) assay, which quantifies intracellular lipid peroxide levels by measuring fluorometric products formed by the reaction of MDA with thiobarbituric acid (TBA). Lipid peroxides were found to be increased in OC cells in which *FZD7* was knocked down compared to control-transduced cells (Fig. 5N, *p* < 0.05), consistent with decreased GPX4 levels. Conversely, lipid peroxides were decreased in OC cells overexpressing *FZD7* (Fig. 5O, *p* < 0.05) supporting that FZD7(+) cells clear these toxic products more effectively, perhaps due to higher levels of GPX4.

GPX4 participates in regulation of intra-cellular redox states by utilizing glutathione (GSH) as the critical antioxidant (24). GSH is synthesized from glutamate-cysteine under the action of glutamate-cysteine ligase (GCL) (24). The cycling of reduced GSH to oxidized glutathione disulfide (GSSG) removes ROS derived from hydrogen peroxide and lipid hydroperoxides through various glutathione peroxidases (GPXs), including GPX4 (24). GSH recycling from GSSH is catalyzed by glutathione reductase (GSR), using NADPH, whose synthesis is regulated by isocitrate dehydrogenase 2 (IDH2) (24). *FZD7* knockdown in OVCAR5 cells, decreased the expression levels of multiple genes in this pathway, including *GPX2, GSS (glutathione synthase), IDH2, GSR, GCLC* and *SLC7A11 (solute carrier family 7 member 11, cystine/glutamate transporter*) (OVCAR5, Fig. 6A and SKOV3, Fig. S7B). Reversely, FZD7 overexpression caused increased expression levels of *GSS, GSR, GCLC and SLC7A11* (OVCAR5, Fig. 6B and SKOV3, Fig S7C), suggesting a significant direct correlation between *FZD7* and glutathione metabolism related genes.

**Figure 6.**
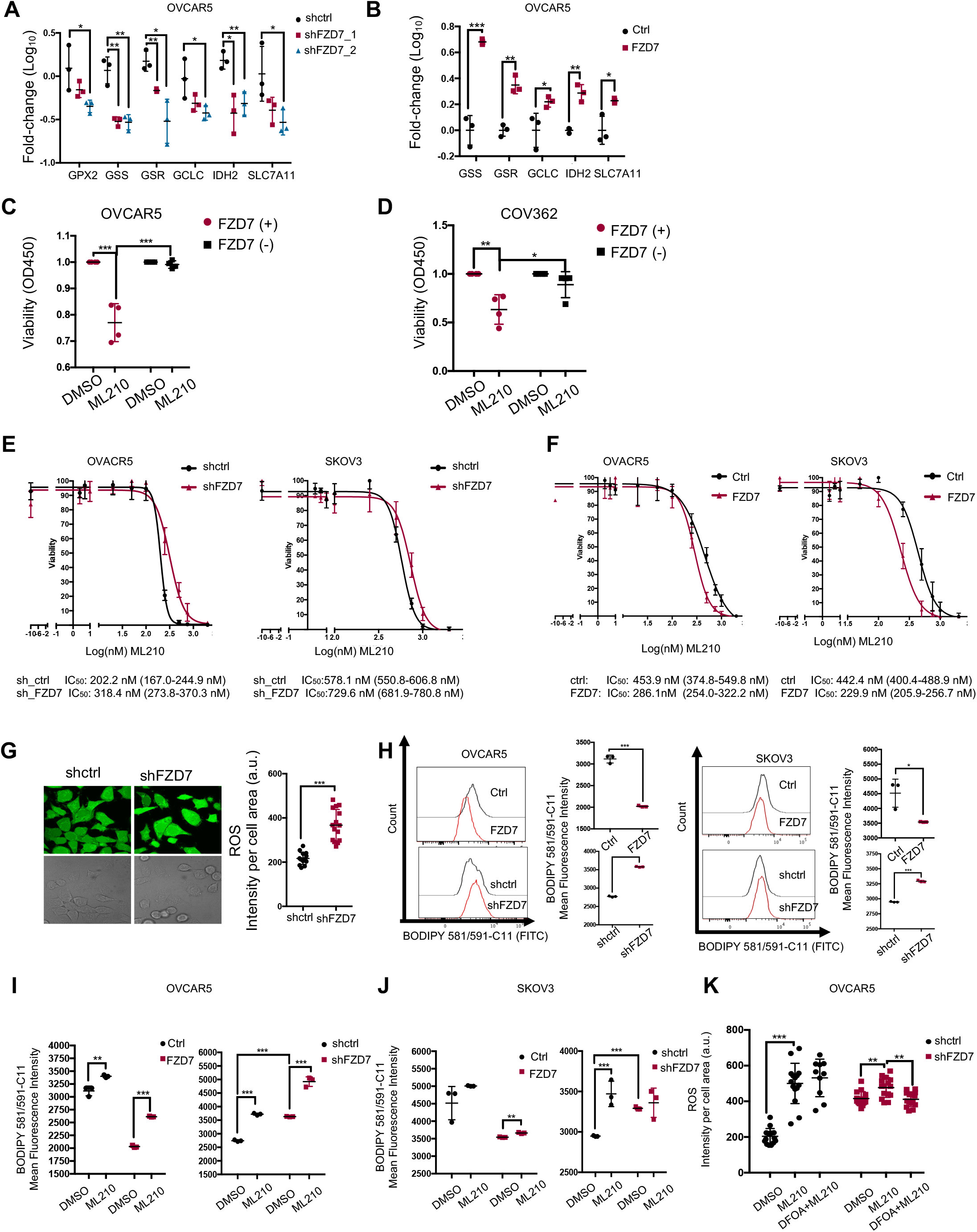
FZD7 marks a cell population susceptible to GPX4 inhibitors. **(A)** Average foldchange (±SD, n = 3) in *mRNA* expression levels of the indicated glutathione metabolism genes in OVCAR5 cells transduced with shRNAs targeting FZD7(shFZD7) vs. cells transduced with control shRNA (A), and in OVCAR5 cells transfected with FZD7-pcDNA3.1 (FZD7) compared with cells transfected with control vector (Ctrl) **(B)** *mRNA* levels were measured by real-time RT-PCR. **(C-D)** Viability of FZD7(+) and FZD7(-) cells sorted by FACS from OVCAR5 and COV362 (D) OC cell lines and treated daily with DMSO or the GPX4 inhibitor ML210 (OVCAR5, 2μM; COV362, 1μM) for 72 hours. Cell viability was determined with a CCK8 assay. Data are presented as average fold-change (± SD, n = 4) of absorbance values relative to control. **(E)** Cell survival curves of OVCAR5 (left) and SKOV3 (right) OC cells transduced with control shRNAs (shctrl) or shRNAs targeting *FZD7* (shFZD7) and treated with GPX4 inhibitor (ML210) for 3 days. Cell viability was measured with the CCK8 assay (n=3-4). ML210 IC_50_ values are shown below figures. **(F)** Survival curves of OVCAR5 (left) and SKOV3 (right) cells transfected with FZD7-pcDNA3.1 or control vector and treated with the GPX4 inhibitor ML210 for 3 days. Cell viability was measured with the CCK8 method. ML210 IC_50_ values are shown below figures (n=3-4). **(G)** Images of intracellular ROS (left), and quantification of intracellular ROS levels (right) in OVCAR5 OC cells transfected with control shRNA (shctrl) or shRNA targeting *FZD7* (shFZD7). ROS levels were measured with a DCF assay. Data are presented as means (± SD) of DCF fluorescence intensity per cell area (n = 15). **(H)** Histograms of fluorescence intensity (left) and mean (± SD, n = 3) (right) fluorescence intensity of BODIPY 581/591-C11 show levels of lipid peroxidation in OVCAR5 (left) and SKOV3 (right) OC cells transfected with empty vector (ctrl), FZD7-pCDNA3.1 (FZD7), control shRNA (shctrl), or shRNAs targeting *FZD7* (shFZD7). BODIPY fluorescence was measured by flow cytometry. **(I, J)** Mean (± SD, n=3) fluorescence intensity of BODIPY 581/591-C11 show effects of ML210 (1 μM for 20 hours) on lipid peroxidation levels in SKOV3 (I) and OVCAR5 (J) cells transfected with empty vector (ctrl), FZD7-pcDNA3.1 (FZD7), control shRNA (shctrl), or shRNA against *FZD7* (shFZD7). BODIPY fluorescence was measured by flow cytometry. **(K)** Intracellular ROS levels in OVCAR5 cells transfected with shctrl and shFZD7 treated with DMSO, ML210 (2μM, 24 hours) and ML210 + DFOA (800nM, 24 hours) were measured by assessing DCFHDA oxidation. Average intensity per cell area (± SD) is shown (n=15). For all comparisons: *P<0.05, **P<0.01, ***P<0.001.

### FZD7 Marks a Cell Population Susceptible to GPX4 Inhibitors

Given the direct correlations observed between FZD7, upregulated in platinum-tolerant cells, and GPX4-mediated cellular redox maintenance, we hypothesized that inhibition of this axis could eliminate resistant cancer cells. Small molecule inhibitors of GPX4 have been shown to increase cellular oxidative stress and to induce ferroptosis, an iron-dependent lipid peroxide-induced cell death (2, 31–33). Thus, we examined the sensitivity of OC cells with high vs. low FZD7 expression levels to GPX4 inhibitors, ML210 and RSL3 (2, 33, 34). FZD7(+) sorted cells were more sensitive to the GPX4 inhibitor ML210 compared to FZD7(-) cells (OVCAR5, Fig. 6C; COV362, Fig. 6D, SKOV3, Fig. S7D). *FZD7* knockdown in OVCAR5 and SKOV3 cells also slightly reduced sensitivity to GPX4 inhibitors (Fig. 6E) compared to shRNA control transduced cells, while *FZD7* overexpression slightly increased sensitivity to ML210 and RSL-3 compared to empty vector-transduced cells (Fig. 6F).

Through its anti-oxidant function, GPX4 protects mitochondria from damage, maintaining normal oxidative phosphorylation. To test whether these processes were altered in FZD7(+) vs. FZD7(-) cells, as a consequence of differential GPX4 expression, oxygen utilization was measured by using fluorescence labeled oxygen uptake assay and the seahorse assays. FZD7 knock down resulted in decreased oxygen uptake (Fig. S7E) and consumption rate (Fig. S7F), supporting the role of this pathway maintaining normal mitochondrial function in OC cells. Additionally, intracellular reactive oxygen species (ROS) levels were quantified by measuring intracellular fluorescence labeled oxidation of cell permeable DCFHDA. *FZD7* KD resulted in increased ROS production (Fig. 6G). As increased ROS levels contribute to oxidation of polyunsaturated lipids, leading to ferroptosis, C11-BODIPY staining was used in cells expressing different levels of FZD7 and/or exposed to GPX4 inhibitors. Oxidation of the polyunsaturated butadienyl portion of the dye in the presence of ROS is reflected in a shift of the fluorescence emission peaks from red to green, and represents a hallmark of ferroptosis. The mean green (FITC) fluorescence intensity caused by oxidized lipids was decreased in cells overexpressing FZD7 and increased in cells in which FZD7 was knocked down (Fig. 6H), consistent with the increased susceptibility to ferroptosis of FZD7(+) cells. ML210-induced increase in fluorescence was higher in SKOV3 and OVCAR5 cells overexpressing FZD7 compared to controls and decreased in cells in which FZD7 was knocked down (Fig. 6I-J). Likewise, baseline and ML210 induced intracellular ROS levels (rescued by the iron chelator DFOA) were higher in OVCAR5 cells transduced with control shRNA compared to cells transduced with shRNA targeting *FZD7* (Fig. 6K). Collectively, the data suggest that cells marked by FZD7 are more susceptible to ferroptosis and could be eliminated from tumors by targeting GPX4.

### FZD7 regulates GPX4 expression and glutathione metabolism by activating canonical b catenin/p63 pathway

As a classical Wnt receptor, FZD7 participates in both canonical β-catenin and non-canonical planar cell polarity (PCP) signaling. One of the known β-catenin targets is the transcription factor *p63*, which is directly transactivated by the TCF/LEF complex (35). Its most common isoform, ΔNTp63, lacking its N-terminus domain, has been implicated in maintaining intracellular redox homeostasis by regulating genes involved in glutathione metabolism, including GPX4 (24). We therefore hypothesized that in platinum-tolerant OC cells, FZD7 could alter glutathione metabolism and protect OC cells from oxidative stress, through activation of β-catenin/TP63 signaling. Thus, we examined *Tp63* expression in platinum tolerant OC models and found upregulated *Tp63* in platinum tolerant PDX (Fig. 7A) and OC cells compared to controls (Fig. S8A). *Tp63* was also significantly overexpressed in FZD7(+) vs. FZD7(-) cells derived from SKOV3, OVCAR5, COV362 and human HGSOC (Fig. 7B). Next, the effects of FZD7 KD and overexpression on TP63 expression levels were examined. FZD7 KD caused decreased *Tp63* expression levels in OVCAR5 (Fig. 7C-D) and SKOV3 cells (Fig. S8B and Fig. 7D), while FZD7 overexpression led to increased Tp63 expression levels in OVCAR5 and SKOV3 cells (Figs. 7E-G). Further, FZD7 KD inhibited β-catenin and Tp63 protein expression levels in SKOV3 and OVCAR5 cells (Fig. 7D), while FZD7 overexpression induced β-catenin and Tp63 protein levels (Fig. 7G). These experimental results were validated by examining the TCGA HGSOC database (36). *FZD7* and *Tp63* expression levels were positively correlated (Fig. 7H, r = 0.278, p < 0.0001) and *Tp63* expression was also positively associated with expression levels of genes related to glutathione metabolism, including GSS, GCLC and SLC7A11 (Fig. S8C, *p* < 0.01). Higher *Tp63* expression levels were also significantly associated with poor overall survival in this patient cohort (Fig. 7I, *p* = 0.0431).

**Figure 7.**
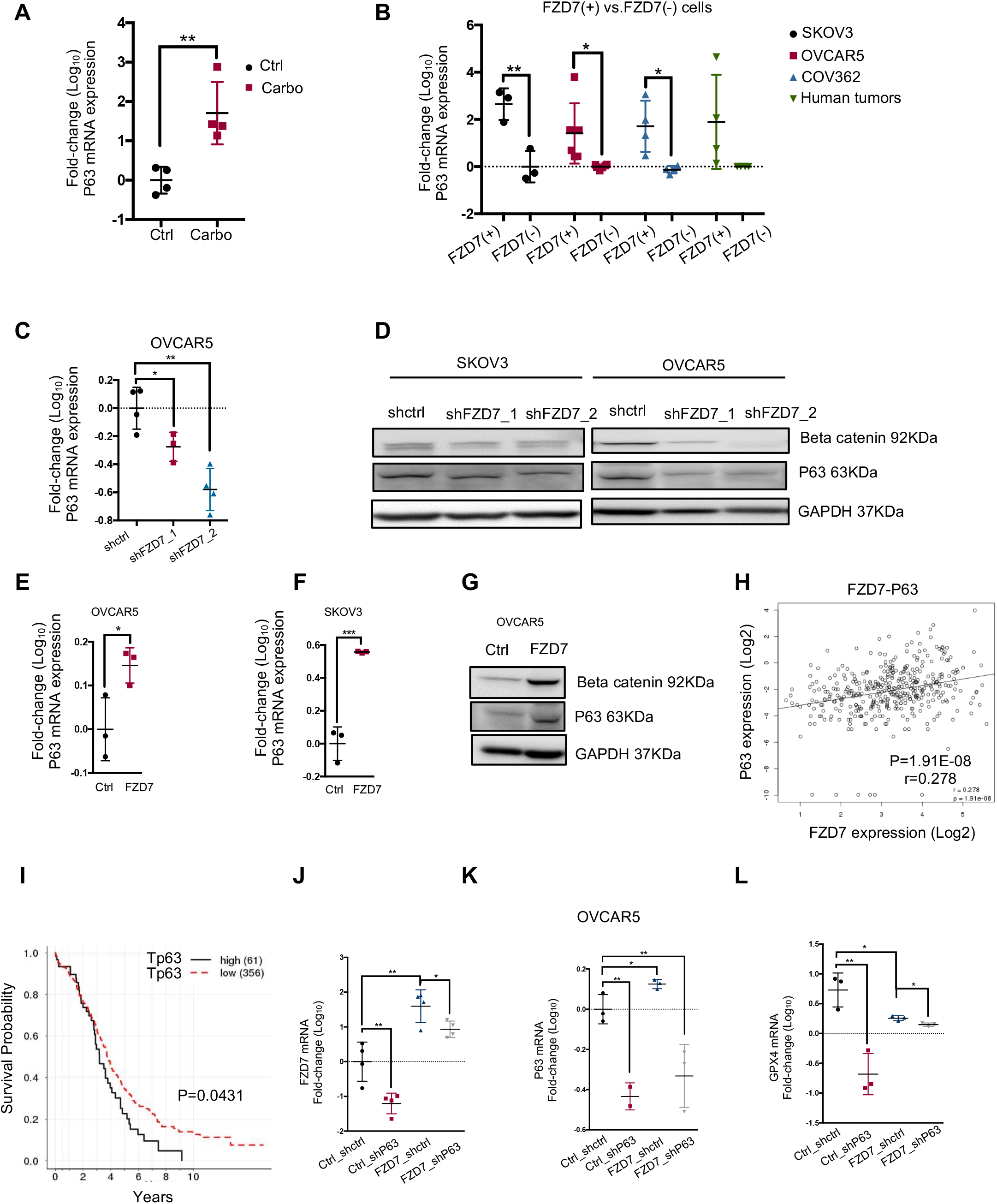
FZD7 regulates GPX4 expression and gluthatione metabolism by activating the canonical b catenin/p63 pathway. **(A)** *P63 mRNA* levels (fold-change ± SD, n=4) measured by real-time RT-PCR in carboplatin tolerant PDX tumors (carbo) compared to control tumors (Ctrl). PDX tumors were produced in mice receiving carboplatin (15 mg/kg, once-a-week for 3 weeks) or PBS treatment. **(B)** *P63* mRNA levels (fold-change ± SD, n=3-6) determined by real-time RT-PCR in FZD7(+) versus FZD7(-) OC cells sorted by FACS from SKOV3, OVCAR5, and COV362 OC cell lines, and cell suspensions from human HGSOC tumors. **(C)** *P63 mRNA* expression levels (fold-change ± SD, n = 4) in OVCAR5 cells transduced with shRNAs targeting *FZD7* (shFZD7) compared with cells transduced with control shRNA (shctrl). *mRNA* levels were measured by realtime RT-PCR. **(D)** Western blot for β-catenin, P63, and GAPDH (loading control) in SKOV3 and OVCAR5 cells transduced with shRNAs targeting*FZD7* (shFZD7) or control shRNA (shctrl) (n=2). **(E, F)** *P63* mRNA levels (fold-change ± SD, n=3) in OVCAR5 (E) and SKOV3 (F) cells transfected with FZD7 expression vector (FZD7) vs. control vector (Ctrl). *mRNA* amounts were measured by real-time RT-PCR. **(G)** Western blot for β-catenin, P63, and GAPDH (loading control) in OVCAR5 cells transfected with control (ctrl) or FZD7-pcDNA3.1 (FZD7) (n = 2). **(H)** Scatter plot shows the correlation between *P63* and *FZD7 mRNA* expression levels in HGSOC tumors (n=419) profiled in the TCGA database. Pearson correlation coefficients and P-values are shown. Transcript *mRNA* levels were measured by RNAseq. **(I)** Kaplan-Meier survival curves for HGSOC patients profiled in the TCGA having high (n=61) or low (n=318) *P63* (T*P63-012*) mRNA expression levels. High or low levels were defined based on statistically determined cutoff point that maximizes absolute value of the standardized two-sample linear rank statistic. **(J-L)** *FZD7*(J), *P63* (K), and *GPX4* (L) mRNA expression levels (fold-change ± SD, n=3-4) in OVCAR5 cells transduced with control shRNAs (Ctrl_shctrl, Ctrl_shP63, FZD7_shctrl) or transfected with FZD7 expression vector and subsequently transduced with shRNA targeting *P63* (FZD7_shP63). *mRNA levels* were determined by real-time RT-PCR. For all comparisons: *P<0.05, **P<0.01, ***P<0.001.

Lastly, to determine whether FZD7 regulates GPX4 expression by altering Tp63 function, the effects of Tp63 knockdown on GPX4 expression in cells expressing high vs. low levels of FZD7 were tested. Overexpression of *FZD7* and knock down of *p63* in OC cells were confirmed at *mRNA* level (OVCAR5, Fig. 7J-K; SKOV3, Fig. S8F-G). *GPX4* upregulation induced by *FZD7* overexpression was abrogated in cells in which *p63* was knocked down (OVCAR5, Fig. 7L; SKOV3, Fig. S8H), supporting that this transcription factor, engaged by β-catenin, downstream of *FZD7*, is an important regulatory node for the glutathione dependent anti-oxidant enzyme. Interestingly, *Tp63* knock down caused reduced expression of *FZD7* in OC cells transfected with both control and FZD7 plasmid vector, indicating a feedback regulatory role of *Tp63* on *FZD7*. In all, our results support the existence of a “persister” cell population, marked by high FZD7 expression, metabolically characterized by increased glutathione dependent anti-oxidant circuits and susceptible to ferroptosis induced by GPX4 inhibitors. A potential mechanism leading to upregulation of GPX4 in platinum-tolerant OC cells marked by FZD7 is engagement of Tp63, trans-activated by β-catenin downstream of this receptor.

## Discussion

Our data support that a platinum-tolerant (“persister”) cancer cell population serving as a reservoir from where resistant tumors emerge, shares common features with CSCs and is characterized by increased expression of the *FZD7* receptor. We demonstrate that the survival of these cells is dependent on an active FZD7-β-catenin-Tp63-GPX4 pathway, which renders these cells susceptible to inducers of ferroptosis. Our findings have several implications.

First, we identified FZD7 as a receptor enriched in platinum-tolerant cancer cells and tumors. FZD7 is a transmembrane receptor which transduces signals involved both the canonical Wnt/β-catenin/TCF and the non-canonical Wnt/PCP signaling pathways (37). Previous data indicated that FZD7 plays essential roles in stem cell biology and in cancer development and progression (38). In breast and hepatocellular carcinoma, FZD7 was linked to oncogenic functions, like cell proliferation, migration, and invasion (39, 40). Here we found that FZD7 marked a population representing ~2-25% cells in cell lines or cells dissociated from human or murine ovarian tumors which were found to be tolerant to platinum. The receptor’s knock down sensitized OC cells to platinum and its overexpression rendered cells more resistant in several OC models. Our group previously identified FZD7 as a receptor facilitating interaction of ovarian CSCs with the tumor niche (41) and our current findings corroborate the link between this receptor and a cancer stemness phenotype. FZD7(+) cells were shown to proliferate more robustly as spheres, to express higher levels of stemness associated TFs, *Sox2* and *Nanog,* and be enriched in a stemness transcriptomic signature. Interestingly, a related receptor, *FZD10,* was recently reported as a marker of resistance to PARP inhibitors (42). Thus, it is likely that activation of the Wnt pathway through activation of one or more of the FZD receptors is linked to resistance to DNA-damaging agents.

Secondly, we report that FZD7 marks a population of cells highly susceptible to ferroptosis. Ferroptosis is a newly described type of cell death that differs from apoptosis and necrosis (43) and is characterized by iron-dependent accumulation of ROS within the cell resulting in increased lipid peroxidation, eventually leading to cell death (34). Cytological changes associated with ferroptotic cell death include cell volume shrinkage and increased mitochondrial membrane density (43). Ferroptosis is dependent on NADPH/H(+), polyunsaturated fatty acid metabolism, and the mevalonate and glutaminolysis metabolic pathways (32) and is inhibited by iron chelation or alpha-tocopherol supplementation (30). Class 1 (system Xc(-) inhibitors) and class 2 (GPX4) inhibitors (43) are two classes of small-molecules that induce ferroptosis (43). Here we observed that FZD7(+) platinum tolerant cells were highly sensitive to ferroptosis induced by small molecules targeting GPX4. Tyrosine kinase inhibitor-tolerant cells and other treatment resistant cancer cells have been recently reported to be sensitive to ferroptosis (2); however, no markers to identify cells prone to ferroptosis have been proposed yet.

Thirdly, we found GPX4 to be significantly upregulated in platinum resistant OC cells, xenografts, PDX models, and importantly, in human ovarian tumors residual after neoadjuvant chemotherapy. GPX4 is a selenoprotein that requires reduced glutathione (GSH) as a cofactor to detoxify lipid peroxides (L-OOHs) by converting them to corresponding alcohols (L-OH). This prevents the buildup of toxic, membrane oriented, lipid reactive oxygen species (L-ROS) generated from oxidation of polyunsaturated fatty acids and cholesterol (44) that, ultimately, kill the cells. In human cells, GPX4 was shown to protect against ferroptosis. In addition, GPX4 also plays a role in maintaining the mitochondrial membrane potential under oxidative stress (29). There are multiple mechanisms to protect cells from ferroptosis, such as increasing GPX4 or GSH levels or by reducing the presence of L-OOHs and/or L-ROSs in the cell membrane. On the other hand, ferroptosis can be induced by mechanisms that reduce GPX4 levels or function, including small molecule inhibitors, reducing GSH levels, or by mechanisms that increase L-OOHs and L-ROSs in the membrane. It had been recently shown that “persister cells” acquire a dependency on GPX4 and loss of GPX4 function induces ferroptosis (2). In colorectal cancer, GPX4 inhibitors were shown to induce ferroptosis by increasing ROS levels and increasing the cellular labile iron pool (34).

Aside from GPX4, other glutathione metabolism related genes, such as *GSH* and *GSR* are also involved in anti-oxidation defense processes that protect cells from free radicals and from chemotherapy attack. Cancer cells with acquired platinum resistance were reported to have increased cellular GSH levels (45). Lower levels of endogenous ROS and higher levels antioxidants and GSH were found in temozolomide (TMZ)-resistant glioblastoma cells. The expression of GSR was also higher in TMZ-resistant compared to sensitive cells and silencing GSR in drug-resistant cells improved sensitivity to TMZ or cisplatin (46). Similar results were reported in clear cell renal carcinoma, where silencing the GSH biosynthesis pathway was shown to trigger ferroptosis (47). Thus, modulation of redox homeostasis by GSH/GSR appears to be an important key modulating sensitivity of cancer cells to chemotherapy (46). Interestingly, in our study, decreased levels of glutathione metabolism-related genes *GSS, GSR, GPX2, IDH* were observed in cells in which FZD7 was knocked down and increased expression of this signature was recorded in cells overexpressing the receptor.

Lastly, our results shed light on a potential mechanism explaining the connection between expression of FZD7 and activation of the glutathione regulatory machinery. A recent study reported that expression of GPX4 and of other genes involved in glutathione metabolism are regulated transcriptionally by *Tp63* (24). Tp63 (along with p53 and p73) belong to the Tp53 family (48, 49). TP63 plays a role in the maintenance of self-renewal and progenitor cell functions in normal epithelial tissues through its by-product ΔNp63, which lacks the N-terminus. ΔNp63 has dominant-negative effects on other isoforms of the p53 family (TAp63, TAp73 and p53) and exerts tumor promoting functions (35). Unlike Tp53, which is inactivated in a majority of human cancers, and especially in OC, p63 is rarely mutated or inactivated in tumors. Importantly, overexpression of Tp63 has been associated with poor survival in OC (48, 49). In our study, exploration of the TCGA database revealed a similar poor prognostic signature associated with high *TP63* expression. Additionally, *Tp63* was shown to regulate expression of *FZD7* and to enhance Wnt signaling in mammary normal and malignant tissue (48). A putative crosstalk between Wnt/b catenin pathway and ΔNp63, the TP63 isoform lacking the N-terminus domain, was reported in skin, hair follicles, mammary glands and limb buds during development (50). We observed a similar direct and strong correlation between TP63 and FZD7 expression levels in OC cells, supporting that expression of this receptor may be regulated by this transcription factor. Furthermore, the expression of GPX4 increased in FZD7 overexpressing cells was downregulated by Tp63 knockdown, supporting this proposed mechanism.

In all, our results propose FZD7 as a new marker identifying OC cell populations likely to survive exposure to platinum and enriched in anti-oxidant response mechanisms. These rare cells responsible for disease relapse after chemotherapy, share stemness features and are susceptible to eradication through ferroptosis. Our data provide compelling evidence that targeting platinum-tolerant FZD7(+) cells with molecules inducing ferroptosis is effective and represents a potential new strategy for a disease of high unmet need.

## Author contributions

YW and DM designed the experiments; YW, GZ, SC, HH, JL, YT and JC performed experiments; GZ, YJ and RD completed bioinformatic analyses; YW, GZ, JL and DM analyzed data; JW and ET provided specimens and edited the manuscript; YW, GZ, HC, YJ, and DM wrote and edited the manuscript; JC provided research reagents and valuable comments; DM provided funding for the study.

## Acknowledgments

This research was supported by funding from the Ovarian Cancer Research Foundation Alliance, the National Cancer Institute (R01-CA224275), and the Diana Princess of Wales endowed Professorship from the Robert H. Comprehensive Cancer Center to D. Matei. Tumor specimens were procured through the Tissue Pathology Core and sequencing was performed in the NUSeq Core supported by NCI CCSG P30 CA060553 awarded to the Robert H Lurie Comprehensive Cancer Center. Flow cytometry analyses were performed in the Northwestern University – Flow Cytometry Core Facility supported by Cancer Center Support Grant NCI CA060553. We thank Dr. Marcus Peter from Northwestern University for valuable comments.

