## Supplemental Material and Methods, Tables and Figures for "Frizzled-7 Identifies Platinum Tolerant Ovarian Cancer Cells Susceptible to Ferroptosis"

#### Supplemental Material (SM)

##### Materials and Methods:

**Cell Culture**: SKOV3 and SKOV3-derived cells were cultured in medium composed of 1:1 combination of MCDB 105 (Sigma Aldrich, Cat# M6395) and Medium 199 (Corning, Cat# 10-060-CV), supplemented with 10% FBS (Fisher Scientific, Cat# 35011CV) and 1% penicillin-streptomycin (Corning, Cat# 30-002-CI). Peo1 and Peo4 cells were cultured in RPMI-1640 with L-glutamine (Corning, Cat#10-040-CV) plus 10% FBS, 1% GlutaMAX (Gibco, Cat# 35050-061), 2mM Sodium Pyruvate (Gibco, Cat# 11360-070), and 1% penicillin-streptomycin. OVCAR5 and OVCAR5-derived cells were maintained in RPMI-1640 with L-glutamine (Corning, Cat# 10-040-CV) plus 10% FBS, 1% GlutaMAX, and 1% penicillin-streptomycin. OVCAR3 and its derived sublines were culture in ATCC-modified RPMI-1640 medium (ATCC, Cat# 30-2001) supplemented with 20% FBS, 1% penicillin-streptomycin, and 0.01 mg/mL recombinant human insulin (Gibco, Cat#12585-014). COV362 cells and its derived sublines were cultured in DMEM with L-glutamine, 4.5g/L glucose, without sodium pyruvate (Corning, Cat#10-017-CV) plus 10% FBS, 1% GlutaMAX, and 1% penicillin-streptomycin. Dissociated cells from OC tumors and xenografts were cultured in ultra-low attachment plates (Corning, Tewksbury, MA, USA) using MammoCult base stem cell medium (Stemcell Technologies, Cat#05620, Cambridge, MA, USA) supplemented with 10% MammoCult proliferation supplements (Stemcell technologies, Cat#05620), 4μg/ml heparin (Stemcell technologies, Cat#07980), 0.48μg/ml hydrocortisone (Stemcell Technologies, Cat#07904), and 1% penicillin-streptomycin.

**Spheroid formation assay.** A serial dilution (500, 1,000, 5,000, 10,000, 50,000 cells) of OVCAR5 cells transfected with shctrl and shFZD7 cells were cultured under non-attachment conditions in 96-well ultra-low attachment plates (Fisher Scientific, Cat#3474) in 100μl

MammoCult base stem cell medium. In other experiments, SKOV3, OVCAR5, COV362 or OVCAR3 cells, including wild type, cisplatin resistant, cells with FZD7 knockdown and FZD7 overexpressing cells, FACS sorted FZD7(+) and FZD7(-) cells derived from OC cell lines, or cells isolated from OVCAR5_shctrl and OVCAR5_shFZD7 xenografts were seeded into 96-well ultra-low attachment plates (1000-20,000 cells/well). Cells were cultured for 7 to 14 days and fresh medium was added every 2 to 3 days. Total numbers of spheroids were counted with an inverted microscope and numbers of viable cells were quantified with a Cell Counting Kit 8 (CCK8) or by measuring intracellular ATP levels with the CellTiter-Glo 3D cell viability assay (Promega, Cat# G9681) following the manufacturer's protocol. Briefly, a volume of CellTiter-Glo 3D Reagent equal to the amount of medium was added into each well, mixed for 30 minutes to induce cell lysis, and incubated for 35 minutes at room temperature to stabilize the luminescent signal. Luminescence was measured by using a microplate reader (SpectraMax GeminiXS, Molecular Devices, San Jose, CA, USA).

**Clonogenic assay.** OC cells (800 to 2,000 cells/well) were seeded in 6 well plates and allowed to attach overnight. Cells received experimental treatments and were cultured for 7–14 days to form colonies. Cell colonies were washed with PBS, fixed with 4% paraformaldehyde and stained with 0.05% crystal violet for counting.

**Patient Derived Xenografts**: Pieces of HGSOC tumors obtained from consenting donors were subcutaneously implanted sc or intra-bursally in female NOD SCID gamma (NSG) mice (The Jackson Laboratory) and allowed to grow to 1-1.5cm diameter over the course of 3-4 months, as previously described (1). Collected PDX tumors were confirmed to be of high-grade serous carcinoma type by histological examination by a board certified pathologist. Pieces of the second generation PDX tumors were subcutaneously re-implanted into NSG mice. Treatment with carboplatin (once-a-week, i.p., 15mg/kg, n=5) or PBS (control, n=5) started when tumors reached 5-7 mm in length, and continued for 6 weeks. Body weights and tumor sizes were measured twice-a-week. Tumors were collected and processed as described.

**Half maximal inhibitory concentration ($IC_{50}$):** FZD7(+) and FZD7 (-) cells derived from SKOV3, OVCAR5, and COV362 cells were sorted by FACS and directly seeded at 10,000 cells/well into 96-well plates. Twenty-four hours after plating, cells were treated with cisplatin (0, 1, 2, 4, 6, 8, 12μM) for 24 hours. Cisplatin was removed (washed off), cells were cultured for 48 or 72 hours and cell viability was measured with a CCK8 kit. In other experiments, 1-5,000 OC cells were seeded in 96-well plates and treated with different concentrations of cisplatin (0, 0.5, 1, 2.5, 5, 10, 25, 50, 75, 100, 250, 500 μM) or carboplatin (0, 0.5, 1, 2.5, 5, 10, 25, 50, 75, 100, 250, 500 μg/ml) and proceeded as described above. To determine the $IC_{50}$ of the glutathione peroxidase inhibitor ML-210, FZD7(+) and FZD7 (-) cells were treated daily with ML-210 (0, 0.25, 0.5, 1, 2μM) or DMSO (0 dose) for 96 hours, and cell viability was determined with a CCK8 kit. In other experiments, 1-5,000 OC cells were seeded in 96-well plates and treated with different concentrations of ML-210 (0, 2.5, 5, 10, 25, 50, 100, 250, 500, 750, 1000, 2000 nM) or RSL-3 (0, 2.5, 5, 10, 25, 50, 100, 250, 500, 750, 1000, 2000 nM) and proceeded as described above.

**Cloning strategy**: The FZD7 coding sequence plus a C-terminal Myc-DDK tag was PCR-amplified from pCMV6-Entry mammalian vector (Origene, cat # RC204167) using forward primer 5'-TATGGATCCATGCGGGACCCCGGCG-3' and reverse primer 5'-TTAGAATTCTTACTTATCGTCGTCATCCTTGTAATCCAGG-3' to introduce BamHI and EcoRI restriction sites. The PCR products were resolved by agarose gel electrophoresis, purified using a QIAquick gel extraction kit (Qiagen, #28704), and digested using BamHI (Thermo Scientific, #FD0054) and EcoRI (Thermo Scientific, #FD0274) endonucleases. Similarly, pcDNA 3.1 empty vector was digested with BamHI and EcoRI to create insertion sites. The DNA insert encoding the FZD7 sequence was then ligated into the pcDNA 3.1 vector. To verify correct insertion, Sanger sequencing was performed using universal T7 and BGH primers flanking the inserted sequence. The sequence of the newly constructed plasmid is given in Supplementary Figure S10.

**RNA extraction and quantitative RT-PCR analysis.** RNA was isolated by using Trizol (Invitrogen, Carlsbad, CA) according to the manufacturer's instructions and quantified with a NanoDrop spectrophotometer (Thermo Scientific). For *mRNA* expression studies, 0.2 to 1 µg of total RNA was reverse-transcribed into cDNA with an iScript cDNA synthesis kit (Bio-Rad, Berkeley, California) according to the manufacturer's instructions. Quantitative real-time PCR analysis was performed using iTaq Universal SYBR Green Supermix (Bio-Rad, Berkeley, California) and a 7900HT real-time PCR instrument (Applied Biosystems, Foster City, CA). The RT-PCR reaction used the following parameters: 94℃ for 10 min, 40 cycles of amplification at 94 ℃ for 15 s and 60 ℃ for 1 min, and an extension step of 7 min at 72℃. Data were normalized using expression of the 18S gene. Relative expression of target genes was calculated using the $2^{-\Delta(\Delta C_T)}$ method where $\Delta C_T = C_{T,\ target} - C_{T,\ 18S}$ and $\Delta(\Delta C_T) = \Delta C_{T,\ stimulated} - \Delta C_{T,\ control}$. Primer sequences (Integrated DNA Technologies, USA) are in Supplemental Table S4. RNA from less than 500,000 FACS sorted cells was isolated with a RNeasy Micro Kit and corresponding protocol (Qiagen, Cat#74004). RNA quantity and quality were determined using a NanoDrop spectrophotometer and a Qubit® 2.0 Fluorometer (Invitrogen). RNA samples were used for real-time RT-PCR as described above.

**Western Blotting:** Protein lysates were prepared using radio immunoprecipitation assay (RIPA) buffer, and protein concentrations were quantified with the Bradford assay (Biorad Protein Assay Reagent, BioRad, Berkeley, CA). Proteins (20μg) were resolved by SDS-PAGE, and electroblotted onto PVDF membranes. The antibodies against P63 (rabbit monoclonal, Cat# ab124762, used at 1:1000), GPX4 (rabbit monoclonal, Cat# ab125066, used at 1:1000) and FZD7 (rabbit polyclonal, Cat# ab64636, used at 1:500) were purchased from Abcam (Cambridge, MA). Mouse monoclonal GAPDH antibody was from Meridian Life Science (Memphis, Tennessee, Cat# H86504M, used at 1:10000). Mouse monoclonal β-catenin antibody was from ECM Biosciences (Versailles, KY, Cat# CM181, used at 1:1000). HRP-conjugated donkey-anti-rabbit polyclonal antibody (Cat#NA9340, used at 1:2000) was purchased from GE Healthcare (Pittsburgh, PA) and HRP-conjugated goat-anti-mouse antibody (Cat#haf007, used at 1:2000) was from R&D System (Minneapolis, MN). Membranes were blocked in TBST containing 5% BSA for an hour, and then incubated with primary antibody at 4℃ overnight and with HRP-conjugated secondary antibody for 1 hour at room temperature. Signal was generated using SuperSignal West Pico PLUS Chemiluminescent Substrate (Thermo Scientific, Cat# 34577) or SuperSignal West Femto Maximum Sensitivity Substrate (Thermo Scientific, Cat# 34095) enhanced chemiluminescent HRP system. Images of protein bands were captured by a luminescent image analyzer with a CCD camera (LAS 3000, Fuji Film) and band intensities were quantified by densitometric analysis using Gel-Pro Analyzer 3.1 software. The antibodies used are included in the main manuscript.

**Immunohistochemistry (IHC):** Sections (5 μm) of paraffin-embedded tissues, or ovarian cancer tissue microarrays were heated at 56℃ for 20 mins and deparaffinized with xylene, followed by re-hydration through decreasing concentration of ethanol (100%, 90%, 70%, 50%, 0%). Antigen retrieval was performed with citrate buffer (10 mM, pH 6.0) for 30 minutes at 95°C as previously described (2). Peroxidase activity was eliminated with 10% hydrogen peroxide (Fisher Scientific, Cat# H324500) for 10 mins, and then tissues were incubated with 0.5% normal goat serum (DAKO, Hamburg, Germany, Cat# K0672) in PBS for 1 hour. Anti-FZD7 (Abcam, Cat# 64636, 1:50), anti-GPX4 (Abcam, Cat# ab125066, 1:250), or rabbit IgG (Santa Cruz, Cat# sc-2027,1:500) were added to tissue sections and incubated overnight at 4°C. This was followed by treatment with avidin-biotin peroxidase reagents of a DAKO Detection Kit (DAKO, Hamburg, Germany, Cat# K0672) and with liquid DAB substrate chromogen (DAKO, Cat# K3467). Sections were counterstained with hematoxylin (Agilent Technologies, Cat# CS700) and cover-slipped. Protein staining was measured as a H-score, which is the product between staining intensity (0 to 3+) and percentage of stained cells (0-100%).

**Cell transfection.** OC cells were seeded, grown to 70% confluence, and then transduced with lentiviral particles containing shRNA in the presence of polybrene (8 μg/ml) for 48 hours. Lentiviral transduction particles containing three shRNAs targeting FZD7 were used (shFZD-1, Cat#TRCN0000008345; shFZD7-2, Cat#TRCN0000008343, and shFZD7-3, Cat#TRCN0000357012). Cells transduced with scrambled shRNA (Mission Lentiviral Transduction Particles, Sigma-Aldrich, St Louis, MO, USA) were used as controls. Stable knockdown of p63 used a pool of lentiviral particles containing shRNA targeting p63 (sc-36161-v, Santa Cruz Biotechnology Inc, Santa Cruz, CA, USA). Control cells were transduced with scrambled shRNA (sc-108080, Santa Cruz Biotechnology Inc.). Transduced cells were selected with puromycin (2 μg/ml for OVCAR5 and SKOV3, and 0.5μg/ml for OVCAR3 cells). FZD7 was cloned into pcDNA3.1 expression vector (see SM). 1μg of the FZD7 expression vector or empty vector was transfected into SKOV3 and OVCAR5 cells using lipofectamine 2000

(Invitrogen, Cat# 11668-019), according to the manufacturer's protocol. Stably transfected cells were selected with 200µg/ml G418 (Geneticin, Thermo Fisher, Cat# 10131027) by using single colony selection strategy. Transiently transfected cells were collected 48 and 72 hours post-transfection.

**RNA Sequencing**: The RNA-seq libraries (n=3 per experimental group) were prepared using the NEBNext Ultra II RNA library prep kit from Illumina (New England Biolabs Inc., Ipswich, MA). *mRNA* was isolated from 1 mg of total RNA and used for first-strand cDNA synthesis. This was followed by second-strand cDNA synthesis, end repair of the cDNA library, dA-tailing of the cDNA library, adaptor ligation, and PCR enrichment. The RNA-seq libraries were checked by using a BioAnalyzer, and then sequenced on an Illumina NextSeq500 system with single-end, 75-bp read length settings. For quality control, raw fastq files were pre-processed using TrimGalore (0.4.4) and cutadapt (1.14) with single-end trimming mode, Phred score cutoff of 20 and minimum sequence length cutoff of 20 bp (3). Trimmed reads were aligned to the ENSEMBL human genome version GRCh38 using STAR (2.5.2)(4) and SAMtools (5). Mapped reads were then counted using HTSeq (6).

**Intracellular reactive oxygen species (ROS).** Intracellular ROS levels were measured by monitoring the oxidation of cell permeable 2′,7′-dichlorofluorescein diacetate (DCFHDA, Sigma-Aldrich) to fluoros-pectrophotomete at excitation and emission wavelengths of 480 and 535 nm, respectively, measuring intracellular hydroxyl, peroxyl and other ROS activity. 150,000 cells cultured in 35 mm glass bottom dish were treated with 1 or 2µM ML210 alone or with 800 nM DFOA for 24 hours. Cell cultures were then treated with 10 µM DCFDA (Abcam, Cat#ab113851) for 15 minutes to detect ROS level through confocal fluorescence microscopy. ROS level was measured as integral fluorescence intensity normalized by the cellular area in a frame (n=15 frames) using ImageJ (https://imagej.nih.gov/ij).

#### Supplemental Tables

**Supplemental Table S1. Generation of platinum tolerant ovarian cancer cells *in vitro*.** $IC_{50}$ of SKOV3, OVCAR3, OVCAR5 and COV362 parental and cisplatin/carboplatin tolerant cells are shown.

| Cell line | $IC_{50}$ Range | Cell Line | $IC_{50}$ |
|---|---|---|---|
| SKOV3 | CDDP: 6.5 µM (5.22 - 8.18 µM) | SKOV3 CDDP_R | CDDP: 11.08 µM (9.73-12.62 µM) |
| | Carbo: 4.16 µg/ml (3.10-5.59 µg/ml) | SKOV3 Carboplatin_R | Carbo: 15.01 µg/ml (10.36-21.74 µg/ml) |
| OVCAR3 | CDDP 0.083 µM (0.00044 to 0.15 µM) | OVCAR3 CDDP_R | CDDP: 0.35 µM (0.23 to 0.52 µM) |
| | Carbo: 2.86 µg/ml (1.58 to 5.17 µg/ml) | OVCAR3_Carboplatin_R | Carbo: 15.2 µg/ml (12.64 to 18.29 µg/ml) |
| OVCAR5 | CDDP:2.59 µM (2.31-2.82 µM) | OVCAR5 CDDP_R | CDDP: 8.38 µM (7.47-9.42 µM) |
| COV362 | CDDP: 3.32 µM (3.05-3.63 µM) | COV362_CDDP_R | CDDP: 8.45 µM (6.58-10.84 µM) |

**Supplemental Table S2. Differentially expressed genes in cisplatin and carboplatin-tolerant SKOV3 cells compared to parental cells** (fold change cutoff is 2, n = 1)

| Function | Refseq | Symbol | CDDP vs. control | Carbo vs. control |
|---|---|---|---|---|
| Membrane protein | NM_000118 | *ENG* | 28.79 | 15.55 |
| | NM_005618 | *DLL1* | 20.98 | 17.50 |
| | NM_001954 | *DDR1* | 13.50 | 12.53 |
| | NM_004448 | *ERBB2* | 10.24 | 8.90 |
| | NM_001018016 | *MUC1* | 9.84 | 6.90 |
| | NM_003507 | *FZD7* | 9.79 | 13.04 |
| | NM_021950 | *MS4A1* | 8.14 | 17.50 |
| | NM_001775 | *CD38* | 8.09 | 17.50 |
| | NM_001719 | *BMP7* | 7.49 | 17.50 |
| | NM_019074 | *DLL4* | 7.49 | 17.50 |
| | NM_000264 | *PTCH1* | 7.49 | 17.50 |
| | NM_001773 | *CD34* | 6.92 | 16.17 |
| | NM_000885 | *ITGA4* | 4.94 | 3.76 |
| | NM_006017 | *PROM1* | 4.50 | 10.50 |
| | NM_017617 | *NOTCH1* | 4.33 | 4.12 |
| | NM_005631 | *SMO* | 4.31 | 10.24 |
| | NM_002659 | *PLAUR* | 4.17 | 3.78 |
| | NM_002203 | *ITGA2* | 3.95 | 2.24 |
| | NM_001627 | *ALCAM* | 2.91 | 2.30 |
| | NM_000610 | *CD44* | 2.81 | 2.87 |
| | NM_000214 | *JAG1* | 2.22 | 3.48 |
| | NM_006288 | *THY1* | 2.15 | 6.20 |
| | NM_004612 | *TGFBR1* | 2.02 | 2.47 |
| Transcriptional factors/activators | NM_024674 | *LIN28A* | 19.90 | 17.50 |
| | NM_003106 | *SOX2* | 10.51 | 8.20 |
| | NM_000116 | *TAZ* | 7.77 | 8.49 |
| | NM_002051 | *GATA3* | 7.49 | 17.50 |
| | NM_000474 | *TWIST1* | 6.79 | 5.31 |
| | NM_057179 | *TWIST2* | 5.13 | 8.65 |
| | NM_005985 | *SNAI1* | 5.09 | 21.17 |
| | NM_004475 | *FLOT2* | 4.88 | 4.41 |
| | NM_014757 | *MAML1* | 4.85 | 8.53 |
| | NM_032682 | *FOXP1* | 3.10 | 6.78 |
| | NM_004392 | *DACH1* | 3.04 | 7.68 |
| | NM_005378 | *MYCN* | 2.98 | 6.95 |
| | NM_014795 | *ZEB2* | 2.79 | 4.12 |
| | NM_001556 | *IKBKB* | 2.74 | 2.70 |
| | NM_030751 | *ZEB1* | 2.17 | 2.22 |

| | | | | |
|---|---|---|---|---|
| Others | NM_178559 | *ABCB5* | 7.80 | 17.50 |
| | NM_000930 | *PLAT* | 5.86 | 6.08 |
| | NM_001101 | *ACTB* | 5.13 | 3.80 |
| | NM_004827 | *ABCG2* | 2.69 | 2.79 |

**Supplemental Table S3: Patients' characteristics**

| | Age | Tumor Type | Stage | Sites |
|---|---|---|---|---|
| Pt #1 | 67 | HGSOC | yPT1, yN0 | ovary |
| Pt #2 | 68 | HGSOC | yPT3c, yN0 | ovary |
| Pt #3 | 74 | HGSOC | yPT3c, yN0 | ovary |
| Pt #4 | 78 | HGSOC | yPT3c, yN0 | ovary |
| Pt #5 | 78 | HGSOC | yPT3 | ovary |
| Pt #6 | 80 | HGSOC | yPT3c | omentum |
| Pt #7 | 61 | HGSOC | yPT3 | ovary |
| Pt #8 | 78 | HGSOC | yPT3a | ovary |
| Pt #9 | 57 | HGSOC | yPT3c, yN0, ypM1 | ovary |
| Pt #10 | 59 | HGSOC | yPT3c | omentum |
| Pt #11 | 78 | HGSOC | ypT3, ypN0 | fallopian tube |
| Pt #12 | 78 | HGSOC | yPT3c | omentum |
| Pt #13 | 50 | HGSOC | yPT3c, yN1 | omentum |
| Pt #14 | 81 | HGSOC | yPT3c | ovary |
| Pt #15 | 70 | HGSOC | yPT3c | ovary |
| Pt #16 | 66 | HGSOC | yPT3a | omentum |
| Pt #17 | 74 | HGSOC | yPT1 | ovary |
| Pt #18 | 60 | HGSOC | yPT3b | omentum |
| Pt #19 | 76 | HGSOC | yPT3c | ovary |
| Pt #20 | 74 | HGSOC | yPT3c | ovary |
| Pt #21 | 72 | HGSOC | ypT3c, ypN0, yM0 | omentum |
| Pt #22 | 78 | HGSOC | ypT3c | ovary |
| Pt #23 | 50 | HGSOC | pT3c | omentum |

**Supplemental Table S4. Primer sequences for selected genes**

| Gene Name | Primer Sequence | |
|---|---|---|
| | Forward (5' to 3') | Reverse (5' to 3') |
| 18S | CGT CTG CCC TAT CAACTTTC | GATGTGGTAGCC GTTTCTC |
| *ALDH1A1* | AGGGGCAGCCATTTCTTCTCA | CACGGGCCTCCTCCACATT |
| *ALDH1A2* | AACAACGAGTGGCAGAACTCAGAGAG | ATCGAAAGGTTTTGATGACGCCCTGC |
| *FZD7* | GCCATCCCGCCGTGTCGTTCTCT | GCACACCATTGCACGTGAATGT |
| *GCLC* | CCCAAACCATCCTACCCTTT | GTGAACCCAGGACAGCCTAA |
| *GPX2* | CGATCCCAAGCTCATCATTT | TAAGGCTCCTCAGGACTGGA |
| *GPX4* | TCAGCAAGATCTGCGTGAAC | GGGGCAGGTCCTTCTCTATC |
| *GSR* | CCAACGTCAAAGGCATCTATGCAG | ATCTTCCGTGAGTCCCACTGTC |
| *GSS* | GACCAGCGTGCCATAGAGAATGA | CATGTGACCTCTCCAGCAGTAGAC |
| *IDH2* | GATGGGAAGACGATTGAGGCTGA | TCAGGAAGTGCTCGTTCAGCTT |
| *Nanog* | AGATGCCTCACACGGAGACT | TTTGCGACACTCTTCTCTGC |
| *Oct4* | CTTCGCAAGCCTCATTTC | GAGAAGGCGAAATCCGAAG |
| *P63* | AGAACGGTGATGGTACGAAGCG | GTACTGCATGAGTTCCAGGGACTC |
| *SLC7A11* | GTTGCGTCTCGAGAGGGTCA | GTCGAGGTCTCCAGAGAAGAGC |
| *Sox2* | TGCTGCCTCTTTAAGACTAGGAC | CCTGGGGCTCAAACTTCTCT |

**Supplemental Figures**
**Supplemental Figure S1**

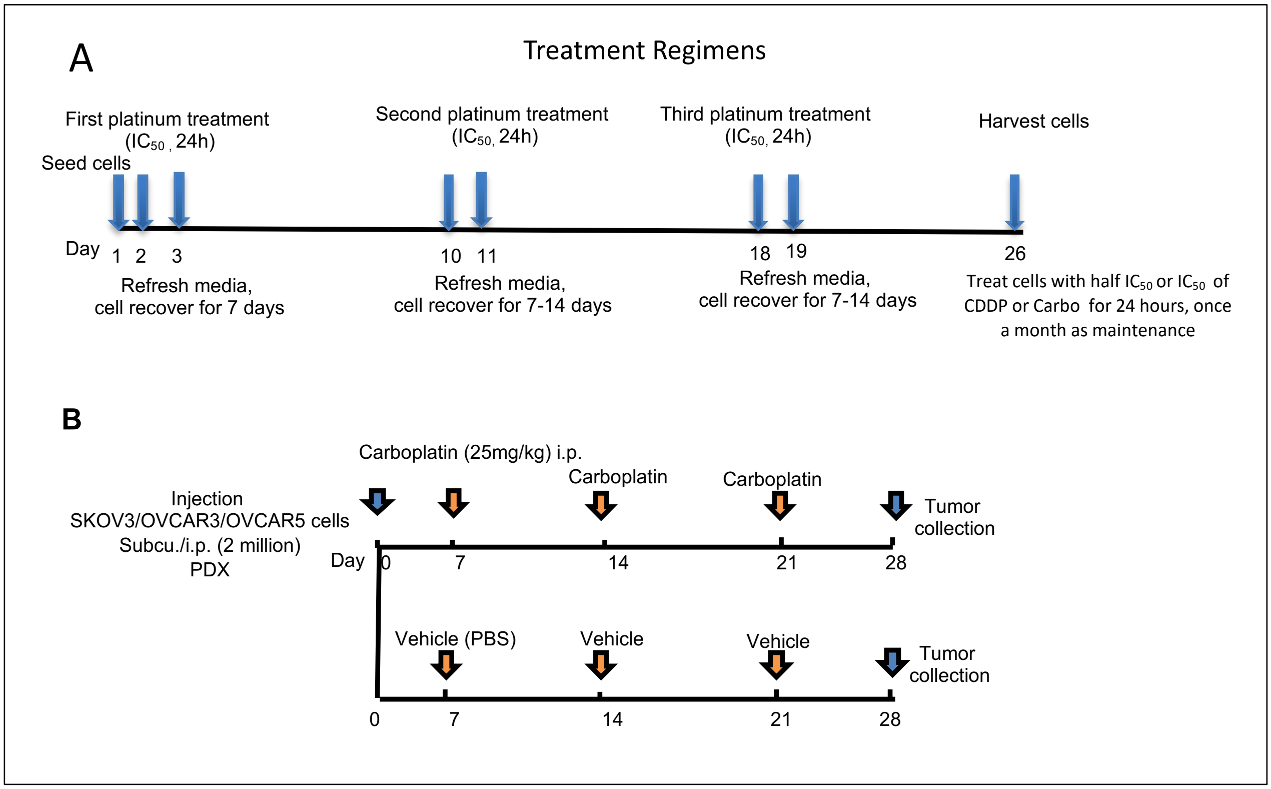


**Supplemental Figure S1 (A)** Schematic diagram describing the development of platinum (cisplatin or carboplatin) tolerant OC cells by using repeated treatment with platinum. **(B)** Schematic diagram describing the development of platinum tolerant ovarian xenografts.

Supplemental Figure S2

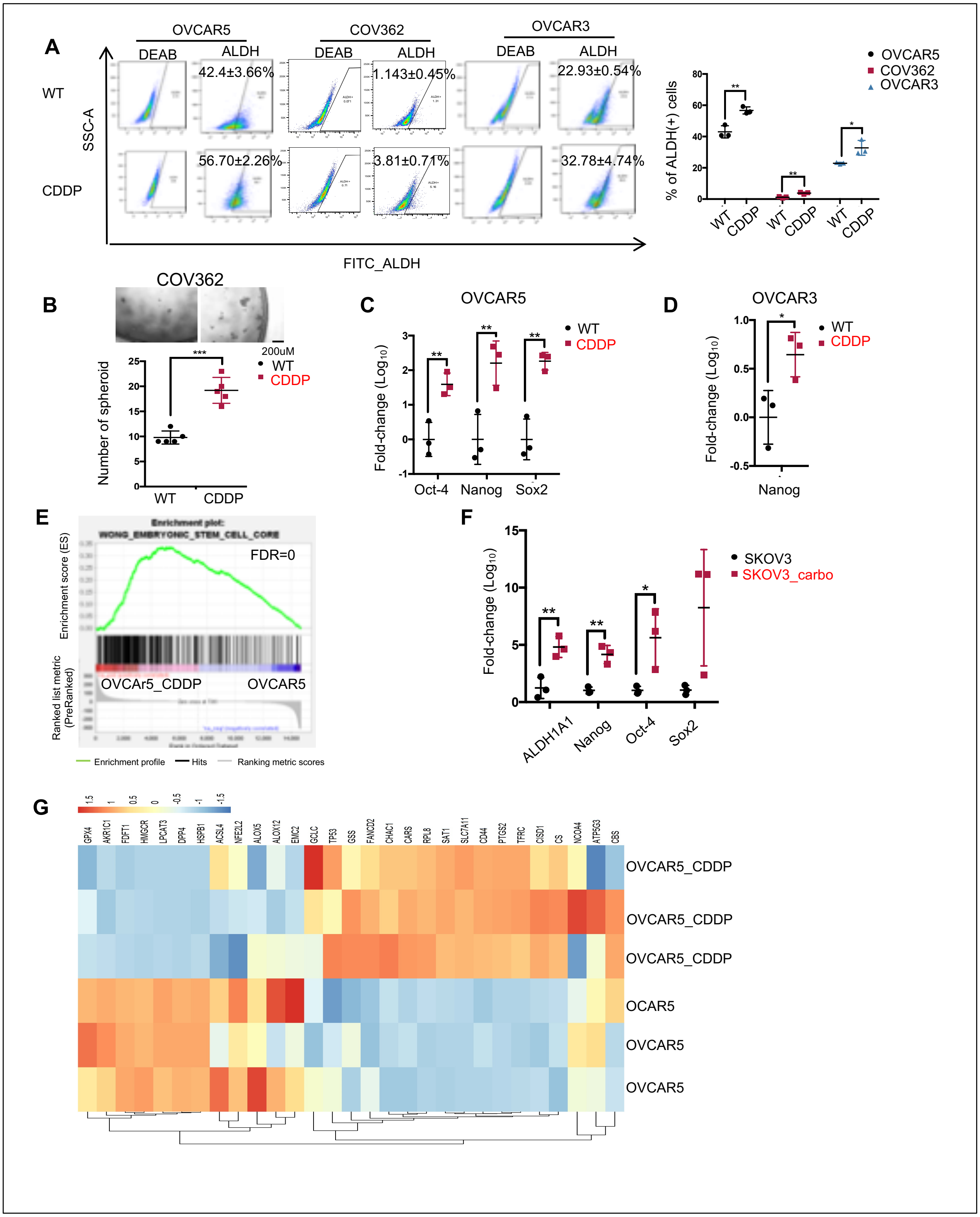

**Supplemental Figure S2.** OVCAR5, COV362 and OVCAR3 cells were repeatedly treated with cisplatin (n = 3 - 4 times). **(A)** Side scatter of FACS shows percentage of ALDH(+) cells in wild type (WT) and cisplatin (CDDP) tolerant cells and mean percentages of ALDH(+) cells (± SD). **(B)** Mean (± SD, n=5) numbers of spheroids generated from 10,000 COV362 CDDP-tolerant cells versus wild type cells (*P< 0.05, **P<0.01, and ***P<0.001). **(C)** Average (± SD, n=3) fold change in mRNA expression levels of stemness associated TFs (*Oct4, Nanog* and *Sox2*) in OVCAR5 CDDP-tolerant vs. parental cells (*P< 0.05, **P<0.01, and ***P<0.001). **(D)** Average (± SD, n=3) fold change of *Nanog* mRNA expression in OVCAR3 CDDP-tolerant vs. wild type cells **(E)** GSEA analysis of Wong Embryonic Stem Cell Core in OVCAR5 CDDP-tolerant vs. wild type cells. Gene list was ranked using signed (from log2FC) likelihood ratio from OVCAR5 CDDP-tolerant vs.parental cells (FDR=0). **(F)** Average fold changes (± SD, n=3) in the expression of *ALDH1A1, Nanog, Oct4* and *Sox2* in carboplatin treated SKOV3 xenografts compared to control tumors (*P< 0.05, **P<0.01, and ***P<0.001). **(G)** Heatmap shows differential expression (FDR<0.05) of ferroptosis related genes between OVCAR5 CDDP-tolerant and parental cells.

Supplemental Figure S3

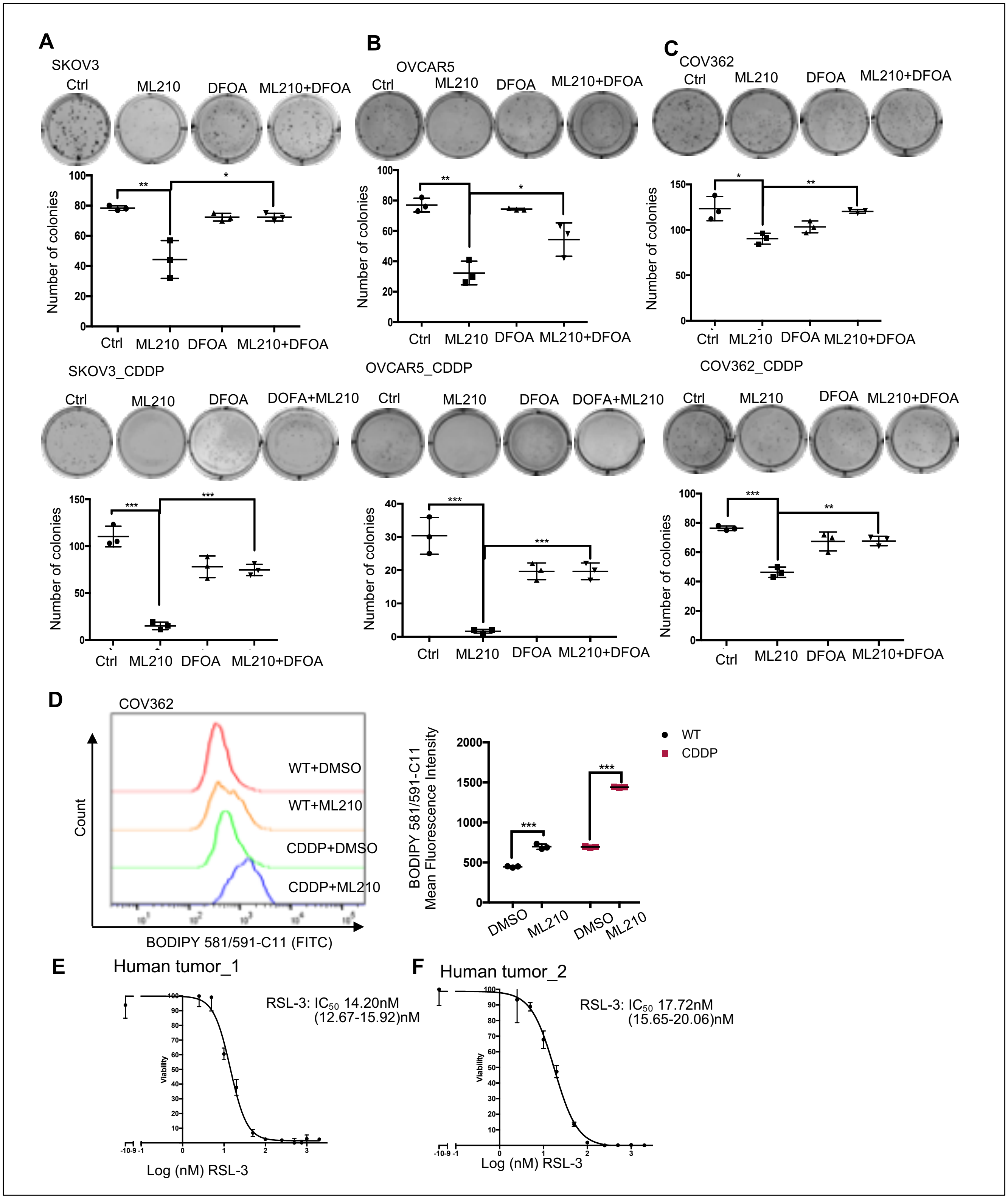

**Supplemental Figure S3. (A)** Colony formation derived from SKOV3, **(B)** OVCAR5, **(C)** COV362 wild type (upper panels) and CDDP-tolerant cells (lower panels) seeded at a density of 1000 cells per well and treated with DMSO, ML210 (SKOV3 and OVCAR5, 1μM; COV362, 2μM), DFOA (800nM) or ML210 and DFOA for 24 hours. Colonies were fixed, crystal violet stained and imaged on Day 14. Average numbers of colonies (± SD) were quantified for each condition (n = 3; *P< 0.05, **P<0.01, and ***P<0.001); **(D)** Lipid peroxidation was assessed in COV362 cells (CDDP-tolerant vs. parental) treated with DMSO and ML210 (2μM for 20 hours) by flow cytometry using BODIPY staining. Histograms are show on the left and mean (± SD, n=3) fluorescence intensities (MFI) of BODIPY 581/591-C11 are shown on the right (***P<0.001). **(E-F)** Cell survival curve of primary platinum-resistant HGSOC tumor cells after treatment with RSL-3 (from 0 to 2000nM) (n=3-4). $IC_{50}$ to RSL-3 are shown.

**Supplemental Figure S4.**

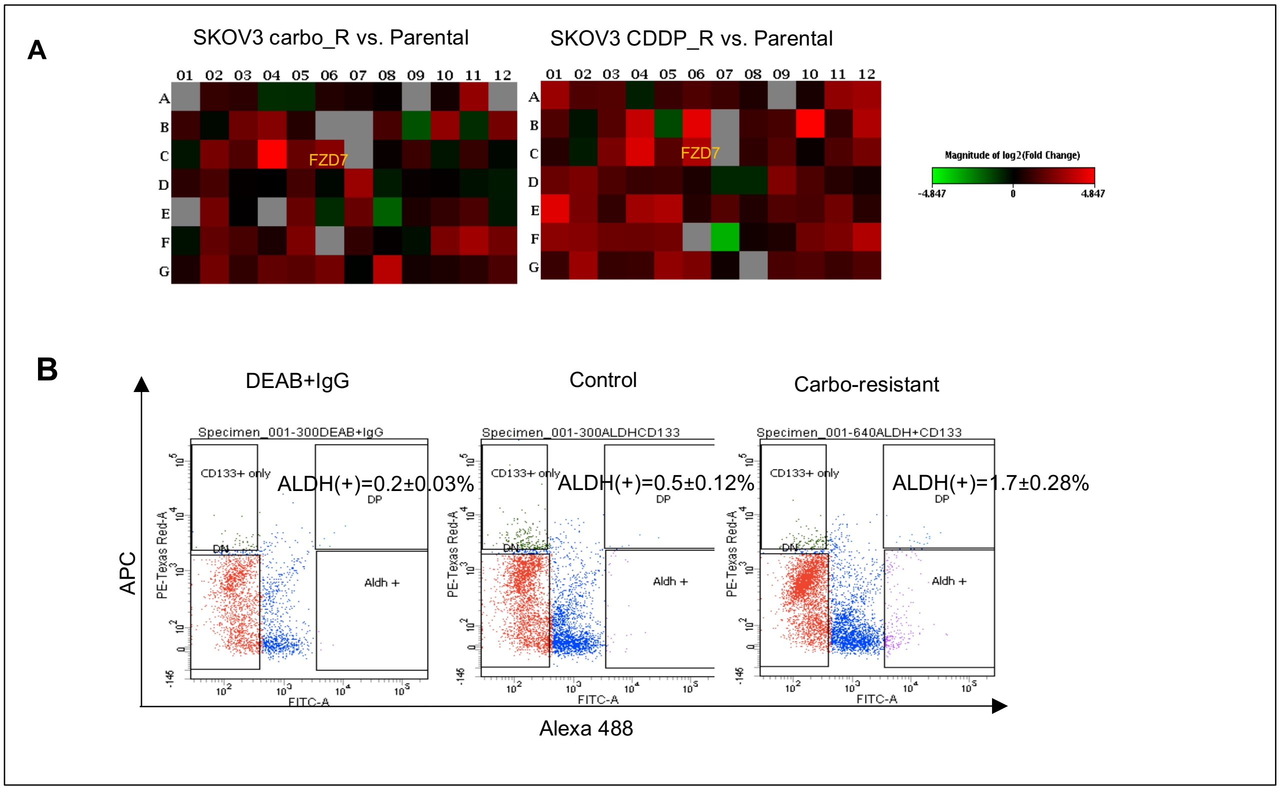


**Supplemental Figure S4. (A)** Heatmap of differentially expressed genes from the cancer stem cell RT-PCR based platform indicates stemness associated genes in the carboplatin (left) or cisplatin (right) tolerant SKOV3 vs parental cells. Red corresponds to upregulated and green to downregulated genes in platinum-tolerant vs. parental cells. **(B)** NSG mice bearing subcutanous PDX ovarian tumors were treated with carboplatin (Sigma) at 15mg/kg, or PBS (n = 5 mice per group). FACS histograms indicate ALDH (+) cells in single cell suspensions derived from PBS or carboplatin-treated PDX tumors; percentages ± SD of ALDH(+)cells were quantified (n=3).

**Supplemental Figure S5.**

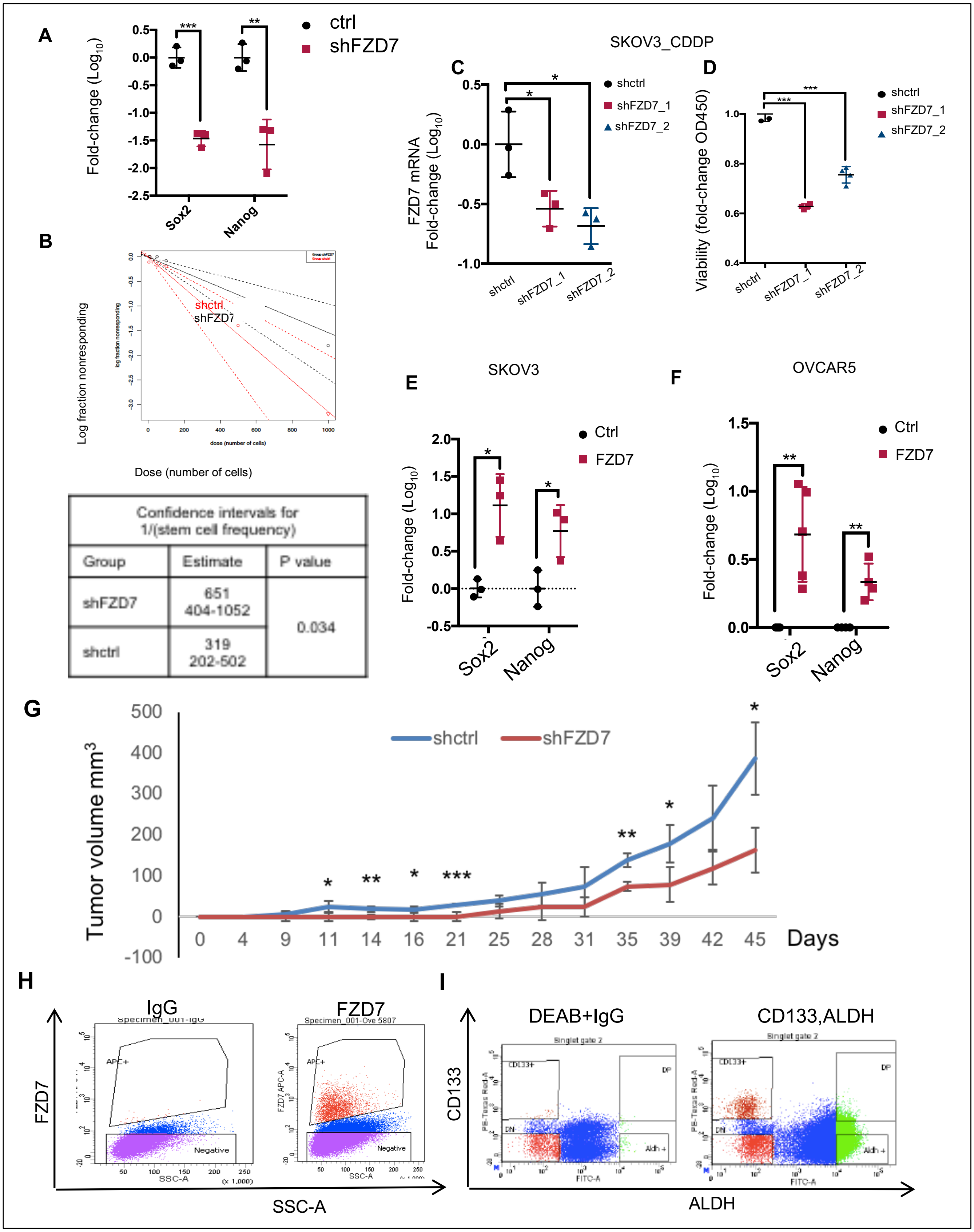


| Confidence intervals for 1/(stem cell frequency) | | |
|---|---|---|
| Group | Estimate | P value |
| shFZD7 | 651<br>404-1052 | 0.034 |
| shctrl | 319<br>202-502 | |

**Supplemental Figure S5. (A)** Mean fold changes (± SD, n=3) of *Sox2* and *Nanog mRNA* expression levels in SKOV3 transduced with shRNA targeting FZD7 (shFZD7) vs. control (shctrl). **(B)** Limited dilution assay used shctrl and shFZD7 transduced OVCAR5 cells. Serially diluted numbers (5, 10, 50, 100, 500 1000) of shctrl or shFZD7 OVCAR5 cells were cultured under spheroid conditions for 7 days (n=10 replicates per condition). Stem cell frequencies were calculated by using the Extreme Limiting Dilution Analysis (http://bioinf.wehi.edu.au/software/elda/) and shown in the table (P= 0.034). **(C)** Average fold changes ($Log_{10}FC$) (± SD, n=3) of *FZD7* mRNA levels in CDDP-tolerant SKOV3 cells transduced with shRNA targeting FZD7 (2 sequences) or control. **(D)** Spheroid formation in CDDP-tolerant SKOV3 cells transduced with shRNA targeting FZD7 (2 sequences) or control. Cell viability was measured and average fold changes (± SD, n=3) are shown (*P< 0.05, **P<0.01, and ***P<0.001). **(E)** Average fold changes (± SD, n=3) of *Sox2* and *Nanog mRNA* levels in SKOV3 and **(F)** OVCAR5 cells transfected with FZD7-pcDNA3.1 vs. empty vector. **(G)** Growth curve of sc xenografts in nude mice generated from 2 X $10^6$ OVCAR5 stably transduced with shRNA targeting FZD7 (shFZD7) vs. control (shctrl). Average tumor volumes (± SD) are shown (n=3/group). **(H)** Gating strategy for FACS sorting of FZD7 (+) and FZD7 (-) OC cells or **(I)** CD133(+) ALDH(+) OCSCs vs. CD133(-) ALDH(-) non-OCSCs from OVCAR5 cells used for RNA sequencing.

### Supplemental Figure S6

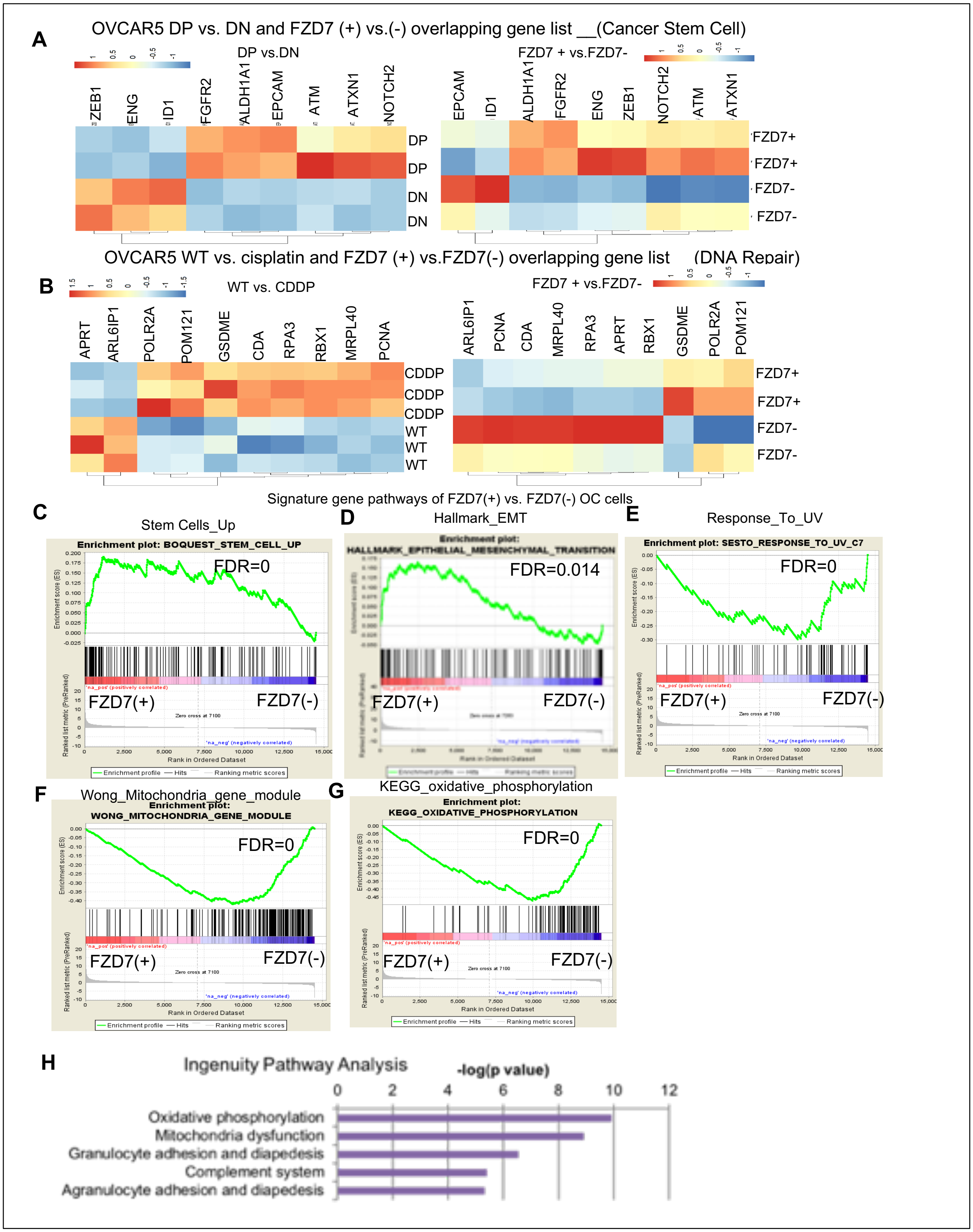

**Supplemental Figure S6. (A)** Heatmap of overlapping differentially expressed genes (FDR<0.05) related to "*Cancer Stem Cells*" pathway between OVCAR5_DP (ALDH+CD133+) OCSCs vs. OVCAR5_DN (ALDH-CD133-) non-OCSCs and OVCAR5_FZD7+ vs. OVCAR5_FZD7- OC cells. **(B)** Heatmap of overlapping differentially expressed genes (FDR<0.05) related to "*DNA Repair*" pathway between OVCAR5 CDDP-tolerant vs. parental and OVCAR5_FZD7(+) vs. OVCAR5_FZD7(-) cells. **(C)** GSEA plots of OVCAR5 derived FZD7(+) vs. FZD7(-) cells related to *Stem Cell UP* (FDR=0), **(D)** *Hallmark Epithelial Meshenchymal Transition* (FDR=0.014), **(E)** *Response to UV gene sets* (FDR=0), **(F)** *Mitochondria Gene_Module* (FDR=0), and **(G)** *KEGG Oxidative Phosphorylation* (FDR=0). **(H)** Ingenuity Pathway Analysis of differentially expressed genes between FZD7(+) vs. FZD7(-) OC cells indicates top canonical pathways enriched in FZD7(+) cells.

#### Supplemental Figure S7

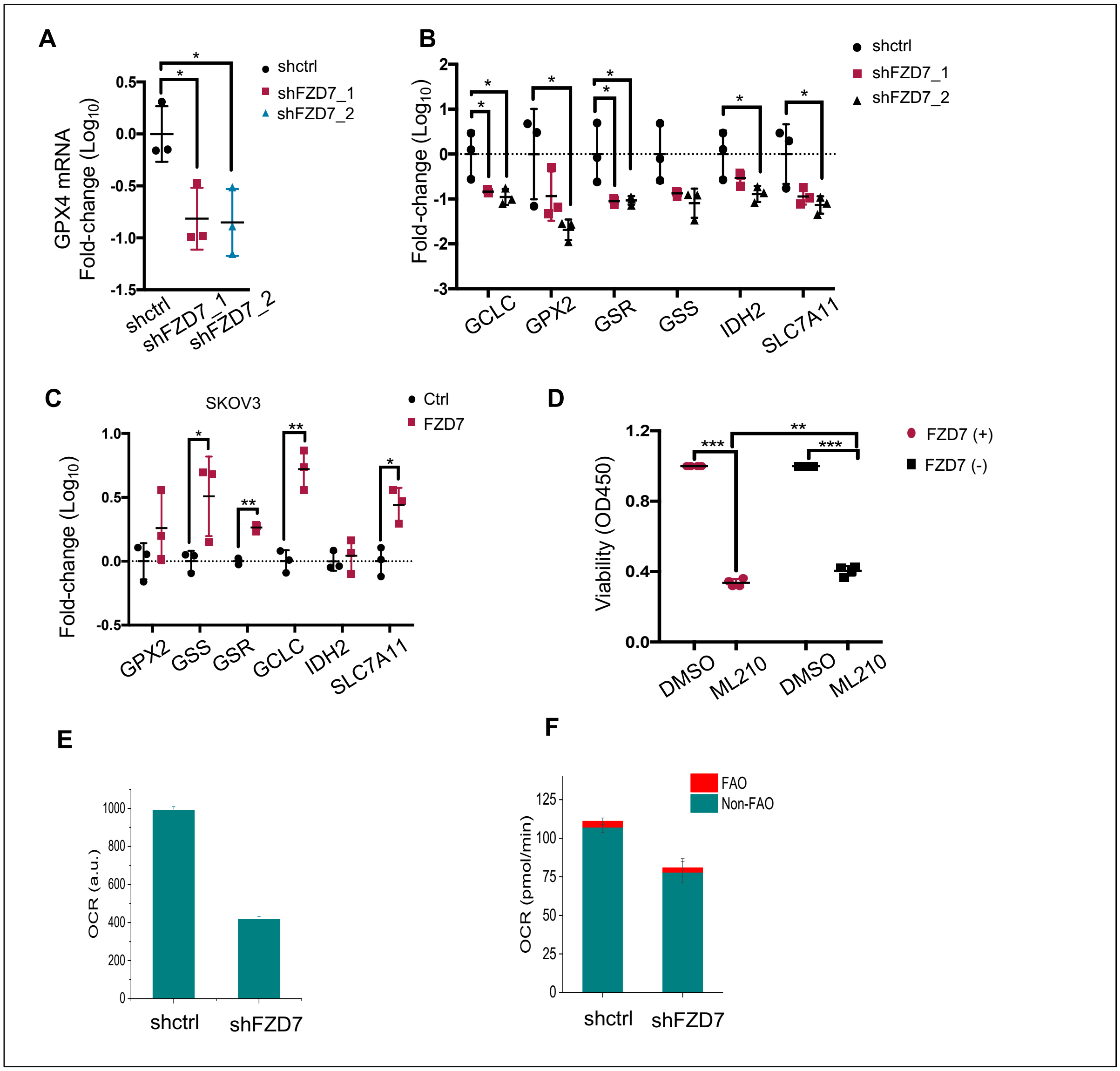


**Supplemental Figure S7. (A)** Mean fold changes (± SD, n=3) of *GPX4 mRNA* expression levels in SKOV3 cells transduced with shRNA targeting FZD7 (shFZD7) vs. control (shctrl) (*P< 0.05, **P<0.01, and ***P<0.001). **(B)** Mean fold change (± SD, n=3) of mRNA expression levels for *GCLC, GPX2, GSR, GSS, IDH2* and *SLC7A11* in SKOV3 transfected with shFZD7 versus SKOV3 transfected with shctrl (*P< 0.05, **P<0.01, and ***P<0.001). **(C)** Average fold change of *GPX2, GSS, GSR, GCLC, IDH2,* and *SLC7A11* mRNA expression (± SD, n=3) in SKOV3 cells transfected with FZD7-pcDNA3.1 vs. empty vector (*P< 0.05, **P<0.01, and ***P<0.001). **(D)** Viability of FZD7(+) and FZD7(-) cells sorted by FACS from SKOV3 cells and treated daily with DMSO or the GPX4 inhibitor ML210 (0.25μM) for 72 hours. Cell viability was determined with a CCK8 assay. Data are presented as average fold-change (± SD, n = 4) of absorbance values relative to control. **(E)** Oxygen consumption rate (OCR) in OVCAR5 cells transduced with shRNA targeting FZD7 (shFZD7) vs. control (shctrl), measured by using the oxygen consumption rate assay kit and **(F)** Seahorse assay (*P< 0.05, **P<0.01, and ***P<0.001).

**Supplemental Figure S8.**

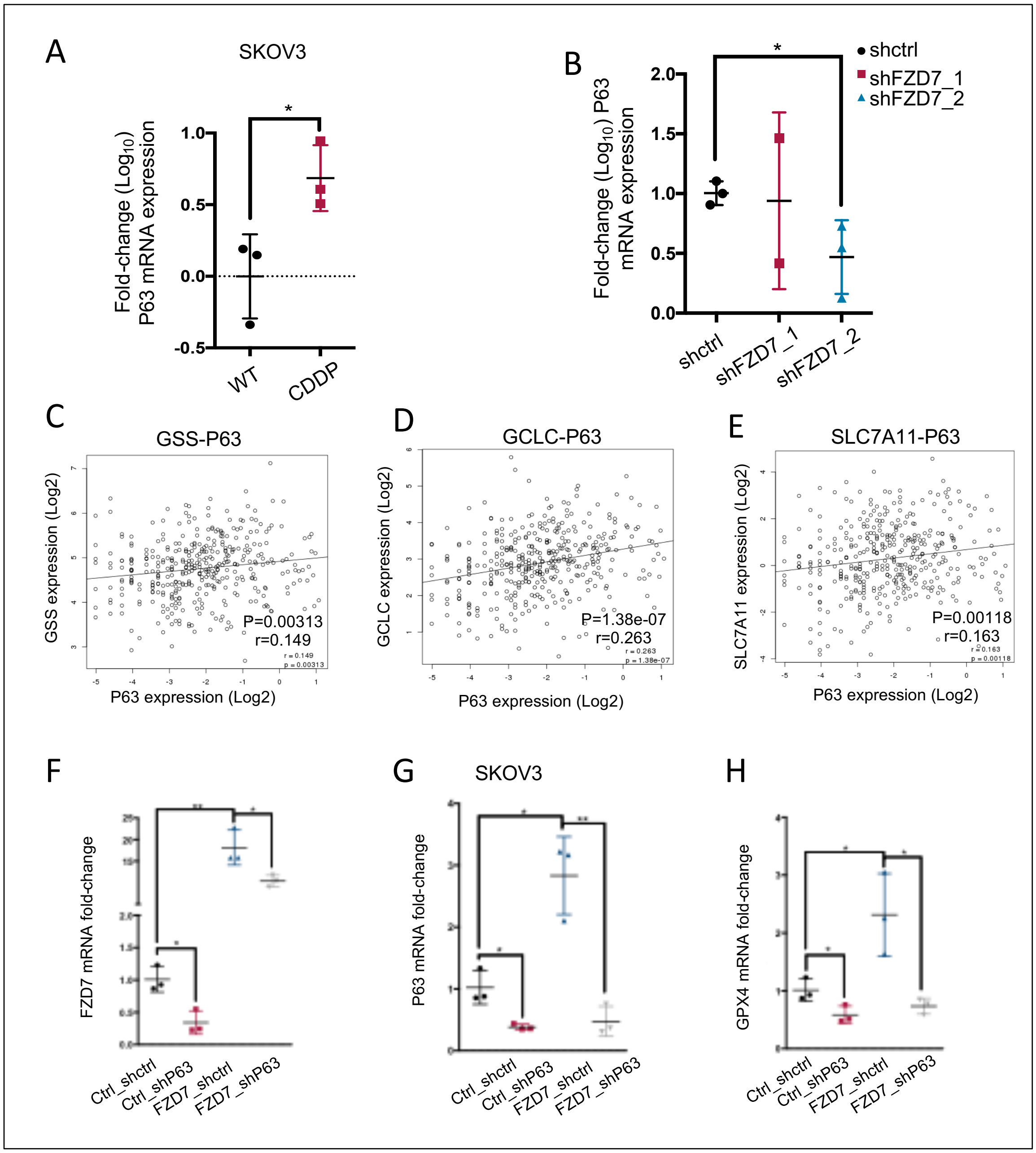


**Supplemental Figure S8. (A)** Mean fold changes (± SD, n=3) of *P63 mRNA* expression levels in SKOV3 CDDP-tolerant vs. parental cells and **(B)** in SKOV3 stably transduced with shFZD7 vs. shctrl (*P< 0.05, **P<0.01, and ***P<0.001). **(C)** Correlations between expression levels of *P63* and gluthatione-related metabolism genes, *GSS* (P=0.00313), **(D)** *GCLC* (P=1.38e-07), and **(E)** *SLC7A11* (P=0.00118), as measured in the TCGA ovarian cancer dataset (n=419). **(F-H)** *FZD7*(F), *P63* (G), and *GPX4* (H) mRNA expression levels (fold-change ± SD, n=3) in SKOV3 cells transduced with control shRNAs (Ctrl_shctrl, Ctrl_shP63, FZD7_shctrl) or transfected with

FZD7 expression vector and subsequently transduced with shRNA targeting *P63* (FZD7_shP63). *mRNA levels* were determined by real-time RT-PCR. For all comparisons: *$P<0.05$, **$P<0.01$, ***$P<0.001$.

.
